# Identification of a new regulation pathway of EGFR and E-cadherin dynamics

**DOI:** 10.1101/2020.07.22.215806

**Authors:** Veronique Proux-Gillardeaux, Tamara Advedissian, Charlotte Perin, Jean-Christophe Gelly, Mireille Viguier, Fredérique Deshayes

## Abstract

EGFR plays key roles in multiple cellular processes such as cell differentiation, cell proliferation, migration and epithelia homeostasis. Phosphorylation of the receptor, intracellular signaling and trafficking are major events regulating EGFR functions. Galectin-7, a soluble lectin expressed in epithelia such as the skin, has been shown to be involved in cell differentiation. Through this study we demonstrate that galectin-7 regulates EGFR function by a direct interaction with its extracellular domain hence modifying its downstream signaling and endocytic pathway. From observations in mice we focused on the molecular mechanisms deciphering the glycosylation dependent interaction between EGFR and galectin-7. Interestingly, we also revealed that galectin-7 is a direct binder of both EGFR and E-cadherin bridging them together. Strikingly this study not only deciphers a new molecular mechanism of EGFR regulation but also points out a novel molecular interaction between EGFR and E-cadherin, two major regulators of the balance between proliferation and differentiation.

**SUMMARY:** EGFR and E-cadherin are known to interact and to regulate epithelial homeostasis. In this study we unravel in the epidermis a new partner and regulator of EGFR which also binds E-cadherin reciprocally bridging their dynamics and functions.

## INTRODUCTION

The Epidermal Growth Factor Receptor (EGFR) is a cell-surface tyrosine kinase receptor that plays a fundamental role regulating cellular metabolism, growth and differentiation by initiating a complex signal transduction cascade (Wee and Wang, 2017). The major EGFR downstream signaling pathways include the mitogen-activated protein kinase (MAPK), the phosphoinositide 3-kinase (PI3K)/Akt, the phospholipase C (PLC), the Janus kinase (JAK), and the signal transducer and activator of transcription (STAT) proteins (Tomas et al., 2014). After EGF binding EGFR is phosphorylated, ubiquitinated and internalized. Two main routes are then possible depending mainly on EGF concentration. In conditions of limited EGF concentration of low EGF, EGFR is predominantly recycled to the plasma membrane rendering it again able to quickly respond to other stimuli whereas at higher concentrations, EGFR is mostly degraded to avoid cell overstimulation (Sigismund et al., 2005). For a long time EGFR was considered to mainly transduce signal at the plasma membrane but studies in the last decade have shown that its endocytosis is not incompatible with the maintenance of signaling (Sousa et al., 2012; Conte and Sigismund, 2016).

Deregulated EGFR signaling and trafficking have been implicated in a substantial percentage of cancers (Yarden, 2001). EGFR is a target of therapy including its inhibition by small molecules interfering with its intracellular kinase domain or by antibodies directed against its extracellular region (Roskoski, 2019). EGFR extracellular domain consists of four subdomains following the signal peptide: I (residues 25 to 189), II (residues 190 to 334), III (residues 335 to 504) and IV (residues 505 to 644) (Roskoski, 2014). These domains are post-translationally modified extensively by sugar unit attachment to Asparagin residues (Asn, N). These glycosylations are major regulators of growth factor binding to EGFR (Azimzadeh Irani et al., 2017).

Glycosylations are also recognized by lectins, proteins with high specificity for these motifs. Among them galectin-7 is a member of the galectin family encompassing soluble lectins with a large variety of ligands and functions. In contrast to certain galectins which are widely expressed in various tissues galectin-7 expression is restricted to pluristratified epithelia such as the epidermis(Madsen et al., 1995; Magnaldo et al., 1998). Furthermore this lectin is observed both in the extracellular compartment and in the cytosol, in mitochondria and even in the nucleus (Advedissian et al., 2017a). Galectin-7 is involved in multiple functions (Advedissian et al., 2017a) such as keratinocyte proliferation and differentiation(Chen et al., 2016), wound healing(Cao et al., 2002, 2003), cell migration and control of cell adhesion through direct interaction with E-cadherin (Advedissian et al., 2017b; Gendronneau et al., 2008, 2015). Galectin-7 is reported in several cancers as a marker but also as an actor in tumor progression (Liu and Rabinovich, 2005; Thijssen et al., 2015).

EGFR is heavily decorated with N-glycans and is involved in cancers notably as a result of aberrant or excessive glycosylation (Manwar Hussain et al., 2018; Hussain et al., 2016), inappropriate *N*-glycosylation often resulting in EGFR dysfunction(Lemmon et al., 2014; Huang et al., 2015). Interestingly, regarding carbohydrate binding, galectin-7 displays preferential binding to internal or terminal LacNAc repeats carried by N-glycan (Rabinovich and Toscano, 2009; Hirabayashi et al., 2002), highly represented on EGFR extracellular domain.

In this study we have chosen to decipher the relationship between EGFR and galectin-7 and the physiological consequences for homeostasis. Hence we unveil that galectin-7 is a direct partner of EGFR *via* interactions through carbohydrate domains. By focusing on downstream signaling and subsequent EGFR endocytosis we demonstrate that galectin-7 is involved in the regulation of EGFR phosphorylation and trafficking. Interestingly we also describe a novel relationship between EGFR galectin-7 and E-cadherin, the major protein involved in adherent junctions already known to be in close proximity with EGFR and to regulate EGFR function (Rübsam et al., 2017). Here we point out for the first time a direct interaction between these three molecules and propose a structural modeling for their interactions.

## MATERIALS AND METHODS

### Cell culture

The HaCaT cell line (Human adult low Calcium high Temperature) were grown in Dulbecco’s Modified Eagle Media (DMEM, Invitrogen) supplemented with 2 mM essential amino acids (Invitrogen), 10 units.ml^−1^ penicillin, 10 μg.ml^−1^ streptomycin (Invitrogen) and 10% foetal bovine serum (FBS) in a 5% CO2 atmosphere at 37°C. Two independent clones with reduced expression of galectin-7 were generated by stable expression of shRNAs as previously described (Advedissian et al., 2017b).

### Animals

Mice were kept on a C57Bl/6 background and housed in a specific pathogen-free animal facility. All experiments were performed on 2 months-old female mice. Animals were handled respecting the French regulations for animal care and wellness, and the Animal Experimentation Ethical Committee Buffon (CEEA-40) approved all mice work.

### Histology and immunostaining

Tissue processing and immunostaining were performed as previously described (Advedissian et al., 2017b). The primary antibodies used are described in supplementary materials.

Nuclei were stained with Hoechst33342 (H3570, Invitrogen) and confocal acquisition was performed using a Leica SP5 microscope.

### Immunofluorescence

Cells were washed with PBS and fixed for 20 min in paraformaldehyde (PFA) 4% at room temperature before being permeabilized for 20min in PBS – 0.025% saponin and then blocked for 30min in PBS – 0.025% saponin – 1% BSA (Bovine Serum Albumin, Sigma-Aldrich). Cells were incubated overnight at 4°C in PBS – 0.025% saponin, 1% BSA containing the primary antibody. The following day, cells were incubated at room temperature in PBS – 0.025% saponin – 1% BSA containing the secondary antibody coupled to a fluorochrome for 1h protected from light. The nuclei were stained with 10μg.ml-1 Hoechst33342 (H357C, Invitrogen). Coverslips were mounted on slides with Fluoromouont-G (0100-01, SouthernBiotech) and visualized using an SP5 confocal scanning Tandem RS (Leica) and analyzed by ImageJ. At least 3 independent experiments have been conducted. Antibodies are detailed in supplementary materials.

### Immuno- and co-immunoprecipitations

Experiments were performed as previously described (Gendronneau et al., 2015). Whole cell extracts were prepared from confluent cells grown on 10 cm tissue culture dishes. For the detection of EGFR ubiquitination, total EGFR Immunoprecipitation was performed with 600μg of proteins from whole cell extracts of HaCaT or shGal-7 cells pretreated with 100ng.ml-1 EGF for the indicated times. The negative control was carried out from a lysate of untreated HaCaT cells. Ubiquitin was revealed with an antibody from Santa Cruz (sc8017, dilution 1/1000).

### *In vitro* binding assay

0.3M of purified proteins (recombinant human E-cadherin 8505 and recombinant human EGFR 344-ER R&D system) were mixed and incubated in 100μl of lysis buffer (25 mM TrisHCl pH 7.5, 100 mM CaCl2, 1 mM EDTA, 1 mM EGTA, 0.5% NP40, 1% TritonX-100, and a protease inhibitor cocktail (11836145001, Roche)) overnight at 4°C under agitation. Then, 60μl of protein-A-Sepharose (P9424, Sigma) was added and samples were incubated 3h at 4°C under agitation. Samples were washed twice with PBS – 0.5% NP40 and twice with PBS before resuspension in Laemmli buffer.

### Western Blot

Proteins (30 to 50μg of total cell lysates) were separated in SDS-PAGE gels and transferred to PVDF membranes (Amersham Hybond-P, GE Healthcare). Membranes were then blocked in PBS-T (PBS – 0.1% Tween 20) supplemented with 5% non-fat milk for 1h at RT and incubated with the primary antibody overnight at 4°C. Immunoblots were visualized using a horseradish peroxidase-conjugated secondary antibody followed by enhanced chemiluminescence detection with an ImageQuant LAS 4000 developer (GE Healthcare).

### Proximity Ligation Assay

The assay was performed with the Duolink^®^ *In Situ* Red Starter kit from Sigma-Aldrich according to the manufacturer’s instructions.

### *In vitro* Wound Healing Assay

Cells were plated in 12-well plates on both sides of a plexiglass insert that was removed once the cells had reached confluence (T0). Cells in the entire well were imaged at T0 and T16 (T= 16 h) and the wound closure was calculated by the difference of the covered area by the cell monolayer between T0 and T16. Images were taken with a Leica MZFLIII system through an Axiocam HRc from Zeiss. Results are mean of three independent experiments performed in triplicate.

### Antibody uptake experiments

Cells were incubated in fresh growth medium containing the anti-E cadherin antibody on ice or at 37°C for different periods of time. Surface-bound antibodies were removed by 1 wash cold PBS then 3 × 5 min acid washes (0.5 M acetic acid, 0.5 M NaCl in PBS) under agitation on ice. Cells were washed with ice-cold PBS++ (PBS +1mM CaCl2 +0.5mM MgCl2), then fixed in 4% paraformaldehyde for 20 min at room temperature and processed for immunofluorescence. The images for quantification were taken with a DMRA2 Leica Microscope. For quantification, the images were background subtracted and cellular regions were identified to measure the total fluorescence intensity using the ImageJ software. Fluorescence intensity measured from cells incubated with E-cadherin antibody 1h on ice were averaged and subtracted from the values measured for the corresponding clones. For each condition tested, three independent experiments were performed and approximately 15 cell groups were analyzed per experiment.

### Cell proliferation assay

Cells were seeded at a density of 5.10^4^ cells per well of 24-well plates and cultured for 2 weeks renewing the medium 3 times a week. Two wells of plated cells per each condition were trypsinized each day the first week and every other day the second week. Total cell numbers were counted in a Malassez cell and expressed as the mean ± standard deviation (SD). Cell number was graphed to obtain the growth curves. Results are mean of three independent experiments performed in duplicate.

### E-cadherin modeling

We modeled the five extracellular domains of human E-cadherin using the mouse structure as template (PDB ID: 3Q2V). Alignment between human and mouse sequences were performed using ORION (Ghouzam et al., 2015, 2016). The percentages of sequence identity and coverage obtained are of 60.8% and 82.3% respectively. The ORION alignment has then been used by MODELLER (Webb and Sali, 2016), resulting in a high-quality model with a DOPE Zscore = −4.27. Z-scores represent the number of standard deviations from the mean of a distribution (generally random) of a given value. The highest the absolute value, the better. Using the same protocol and the same template, we modeled the third domain of E-cadherin extracellular region which has a sequence identity of 76.7% with the template with a coverage of 100%. The resulting model has a DOPE Z-score of −2.5.

### *In silico* study of E-cadherin/Galectin-7 interaction

The study of interaction between human E-cadherin model and galectin-7 crystal structure (PDB ID: 1BKZ) was performed through docking simulations using MEGADOCK 4) (Ohue et al., 2014). 54,000 poses were generated with 3 predictions per each rotation and default scoring function, resulting in more than 162,000 docking experiments.

To model the interaction of galectin-7 with 2 E-cadherin molecules, the first E-cadherin was blocked to avoid E-cadherin/E-cadherin interactions. Galectin-7 is a highly symmetrical homodimer except for the seven first residues of the N-ter extremity which are in open conformation in chain B and close conformation in chain A. Because these residues are important for E-cadherin/galectin-7 interaction and are inherently flexible, we replaced chain A by a duplication of chain B. Expect for this small region both chains are very close to each other (global RMSD of 0.3Å). Each E-cadherin molecule forms the same contacts with galectin-7, resulting in a set of consensus residues that might be responsible for the interaction.

### Modeling of full E-cadherin/Galectin-7 complex

Construction of the full E-cadherin-galectin-7 complex has been built using the galectin-7 docked with two copies of E-cadherin ectodomain 3. E-cadherin model of the five ectodomains (PDB: 3QV2) have then been aligned on both copies of ectodomain 3, resulting in a galectin-7 molecule interacting with two complete E-cadherin. Finally, the full complex has been manually positioned to membrane model containing phosphatidylcholines and phosphatidylserines. Lipids lipid content follow the protocol of Arkhipov et al.(Arkhipov et al., 2013): 30% POPS and 70% POPC in intracellular side and 100% POPC in extracellular side.

## RESULTS

### Galectin-7 depletion impairs epidermis differentiation

To decipher the impact of galectin-7 depletion *in vivo* we used galectin-7 null mice generated in our laboratory (Gendronneau et al., 2008). Refined observations of tail skin of both wild-type and gal7^−/−^ mice revealed the thickening of the epidermis with an accumulation of round cells at the basal layer in absence of galectin-7 (figure 1A). We thus studied the distribution of keratin 14 (K14) and keratin 10 (K10) which are respectively markers of basal undifferentiated and differentiated keratinocytes of the epidermis upper layers (Magnaldo et al., 1998). Mice deficient for galectin-7 instead of having a single layer of basal cells observed as in wild-type mice, exhibit two or even more layers of cells expressing K14 (figure 1B). In addition, a large number of cells with a double K10 / K14 labeling can be observed in comparison to the control. To assess these results cultured cell model, we used HaCaT cell line, an immortalized keratinocyte human cell line, and generated two HaCaT stable clones with a highly reduced expression of galectin-7 (ShGal7) thanks to the expression of an shRNA targeting galectin-7 mRNA as previously described [4] (Figure S1A). We performed RT-qPCR unravelling that galectin-7 depletion induced a strong reduction of K10 mRNA expression, while no modification of K14 could be detected (figure S1B). Consistently, addition of soluble galectin-7 on HaCaT cells led to the increase of K10 transcription (figure S1C).

**Figure 1:**
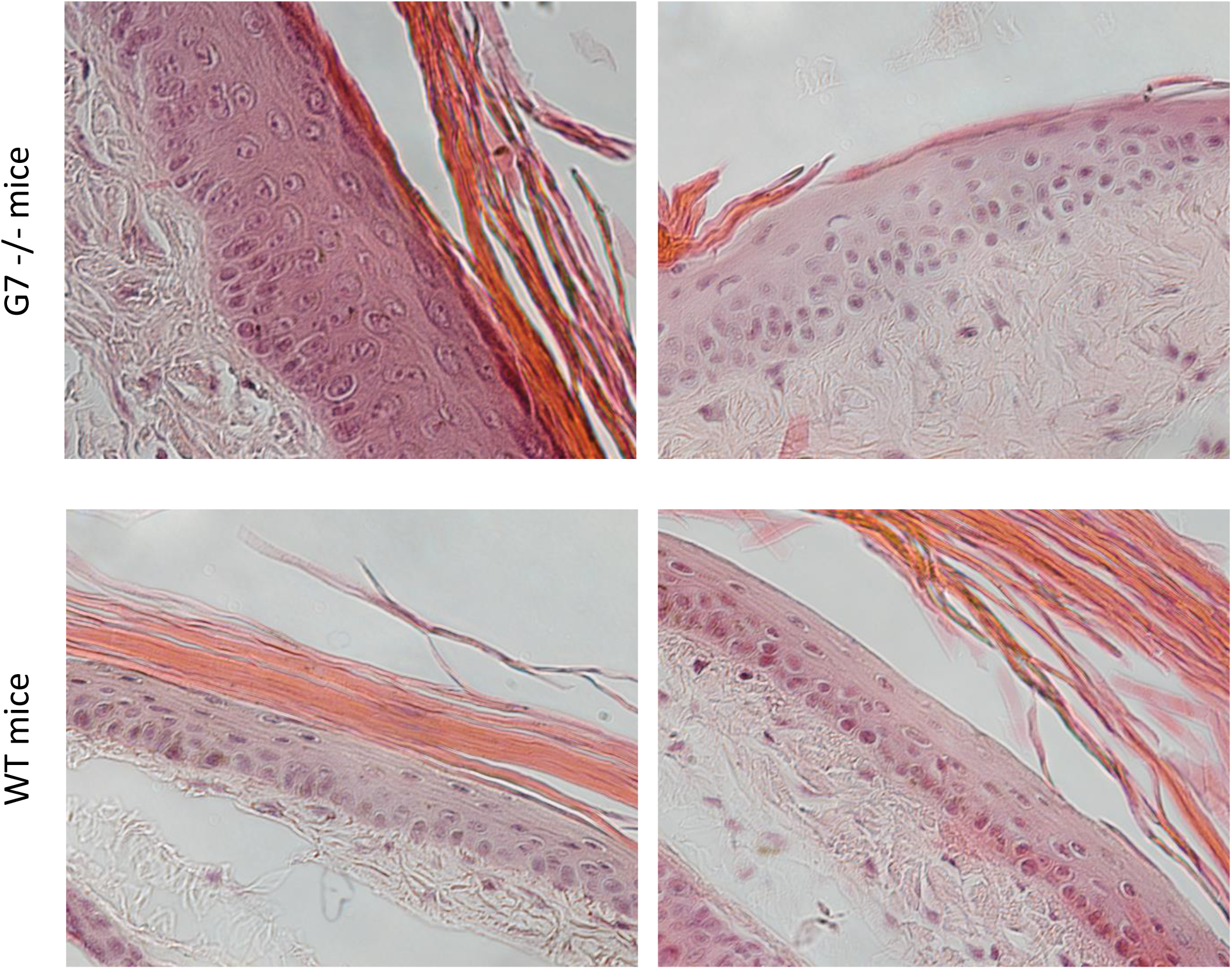
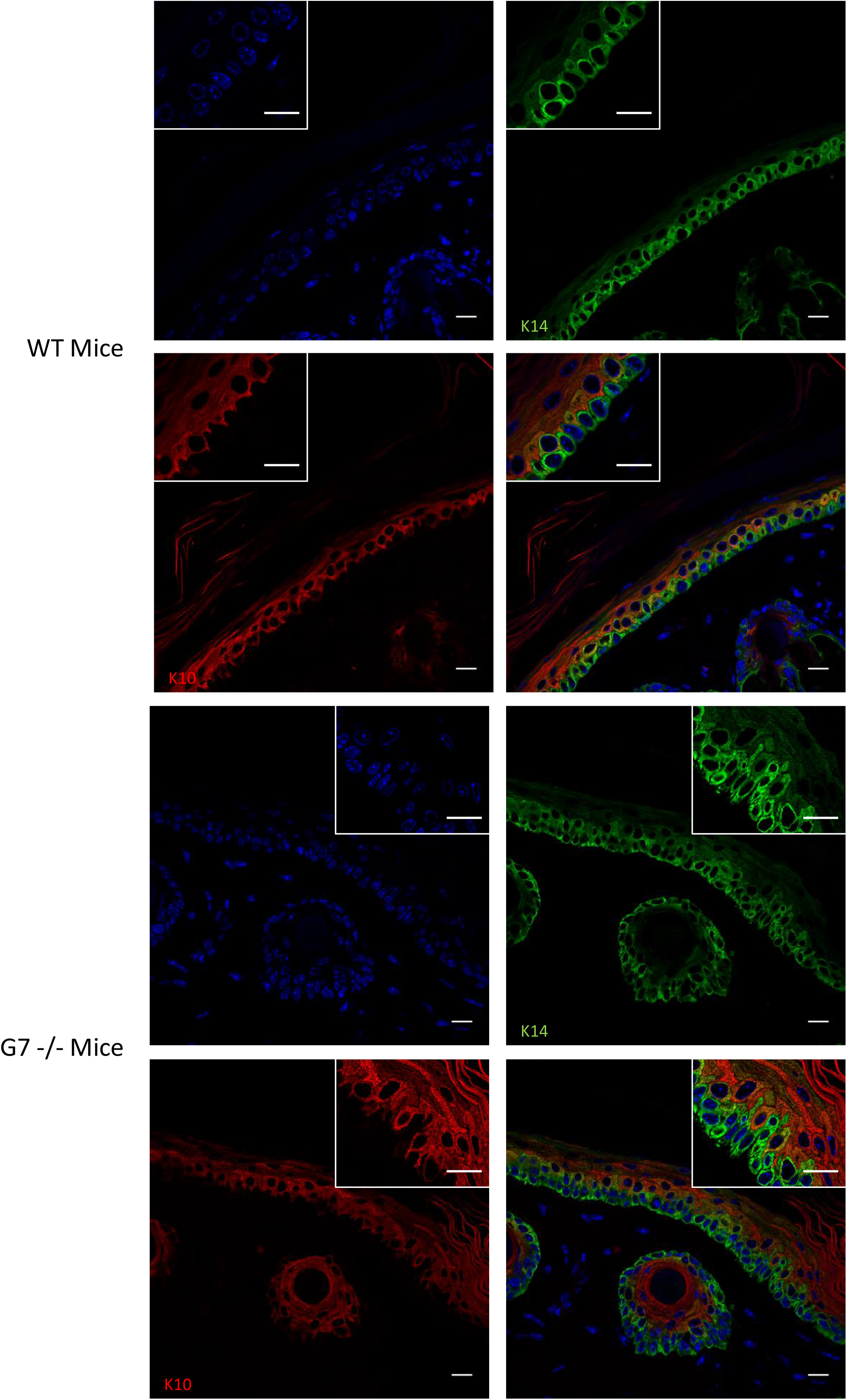
Absence of galectin-7 impairs skin differentiation. A/ Representative staining of wild type and Gal7−/− mice tail with Hematoxylin/Eosin. A thickening of the epidermis is well observable in Gal7−/− mice. Magnification 40x. B/ Representative immunostaining of keratin 14 (green) and keratin 10 (red) in WT and gal7 KO mice tail epidermis showing localization of these two proteins. Scale bar = 15μm

EGFR is known to be a regulator between cell proliferation and cell differentiation. Indeed, it negatively regulates cell differentiation hence decreasing K10 expression. Relying on our observations in mice we studied the effect of the addition of EGF at 100ng/ml for 16 hours on HaCaT cells and studied the level of K10 transcripts. As expected (Chen et al., 2016) EGF induces a significant decrease of K10 transcription in HaCaT cells, having a profile similar to the one observed in absence of galectin-7, whereas only a not significant tendency is observed in the cells without galectin-7 probably due to their already weak level of expression (figure S1C). These observations led us to conclude that galectin-7 is involved in keratinocytes differentiation and appears to have an antagonist effect compare to EGFR signaling. Indeed, the presence of galectin-7 favors cell mechanisms resulting in K10 expression and cell differentiation both *in vivo* and in cultured cells.

### Galectin-7 directly interacts with EGFR through glycosylation of its extracellular domain

The involvement of both galectin-7 and EGF in keratinocytes differentiation correlated with our unpublished preliminary data of proteomic studies suggesting a potential physical interaction between galectin-7 and EGFR. In order to address this question, we performed immunoprecipitation of galectin-7 followed by western blots revealing that galectin-7 does co-precipitate EGFR in HaCaT cells (Figure 2A). To better define the relationship between EGFR and galectin-7, we performed *in vitro* pull-down experiments with purified proteins. A recombinant chimeric protein produced in mouse cells encompassing the EGFR ectodomain (from Met1 to Ser645) fused to an Fc fragment thus called EGFR-Fc was used as a bait, immobilized on beads and incubated with recombinant human galectin-7 (rGal7). As it can be observed on Figure 2C galectin-7 has been retained by EGFR-FC, demonstrating a direct interaction between these two proteins implicating at least the EGFR extracellular domain.

**Figure 2:**
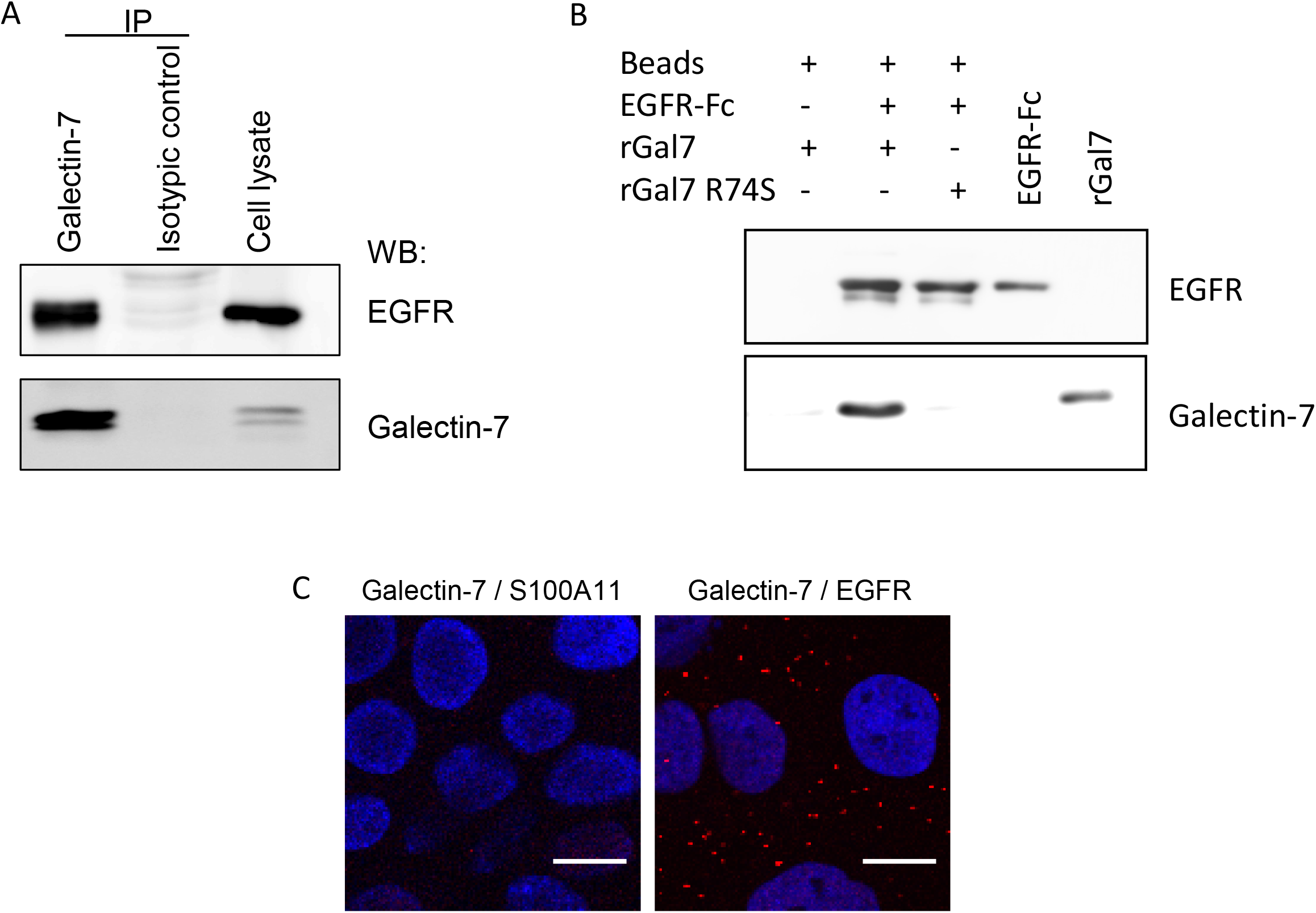
Galectin-7 interacts and colocalizes with EGFR.A/ Co-immunoprecipitation experiments indicate that galectin-7 is a partner of EGFR. Images shown are representative for images taken from distinct western blots. Galectin-7 directly interacts with the extracellular domain of E-cadherin independently of glycosylation motifs. B/ In *vitro* binding assays were performed using recombinant wild type human galectin-7 (rGal7); CRD mutated human galectin-7 (R74S) and extracellular domain of human EGFR fused to human IgG1 Fc fragment (EGFR-Fc). WT galectin-7 (rGal7) precipitated with EGFR-Fc on the contrary of mutated galectin-7 (R74S) C/ Confocal images of Proximity Ligation Assays confirming that galectin-7 is in close proximity with EGFR in cellular context. Galectin-7 – S100A11 pairs were used as negative controls. At least 3 independent experiments were conducted.

In order to further characterize this interaction *in vivo*, we then performed a proximity ligation assay (PLA) allowing the visualization of interactions between two proteins in close proximity (less than 40 nm) with the appearance of red fluorescent dots at the location of their interaction. As observed in figure 2c, galectin-7 and EGFR do interact under these conditions. Indeed, red dots are readily observed at the plasma membrane and sparsely inside cells.

Galectins are widely described as β-galactoside-binding proteins but they can also establish direct protein-protein interactions (Advedissian et al., 2017b). As EGFR is known to be heavily glycosylated, we thus explored the importance of these glycosylation motifs for galectin-7 interaction by using a mutated form of galectin-7 with a R74S substitution in the carbohydrate recognition domain (CRD) which cannot bind to glycosylated motifs. As shown on figure 2C, this form of galectin-7 is not coprecipitated with EGFR-FC *in vitro*. Hence this defective binding with the CRD-mutant suggests a glycosylation-dependent interaction between galectin-7 and the EGFR extracellular domain.

Altogether these data prompted us to conclude that galectin-7 and EGFR physically interact in keratinocytes. We then wondered if this interaction could regulate EGFR activity.

### Galectin-7 depletion favors EGFR phosphorylation and impact its downstream signaling

EGF-binding to EGFR induces multiple events including receptor dimerization, activation of intrinsic tyrosine kinase activity and autophosphorylation. We thus examined EGFR phosphorylation level in both control and ShGal7 clones. After an overnight starvation to remove growth factors, we treated cells with 100ng/ml of EGF or diluent during 15min and cell extracts have been submitted to Western blot analysis. Remarkably, in absence of exogenously added EGF, EGFR displays a basal activation levels with a threefold phosphorylation increase in ShGal7 cell line in comparison to HaCaT cells (figure 3A, B). This is not due to a higher amount of EGFR present at the plasma membrane as assessed by a cell surface biotinylation assay (data not shown). Thus galectin-7 plays a major role in the regulation of EGFR phosphorylation basal level. EGFR possesses multiple downstream targets such as Src, Akt or ERK. Hence, we compared the activation of these different pathways in presence or in absence of galectin-7. We did not observe any differences concerning Src pathway in spite of testing tyrosine 416 phosphorylation site (figure S2A) but we observed an increased phosphorylation on ERK and Akt in absence of galectin-7 (figure 3A, B). Thus galectin-7 restrains basal level of signaling pathways depending on EGFR phosphorylation.

**Figure 3:**
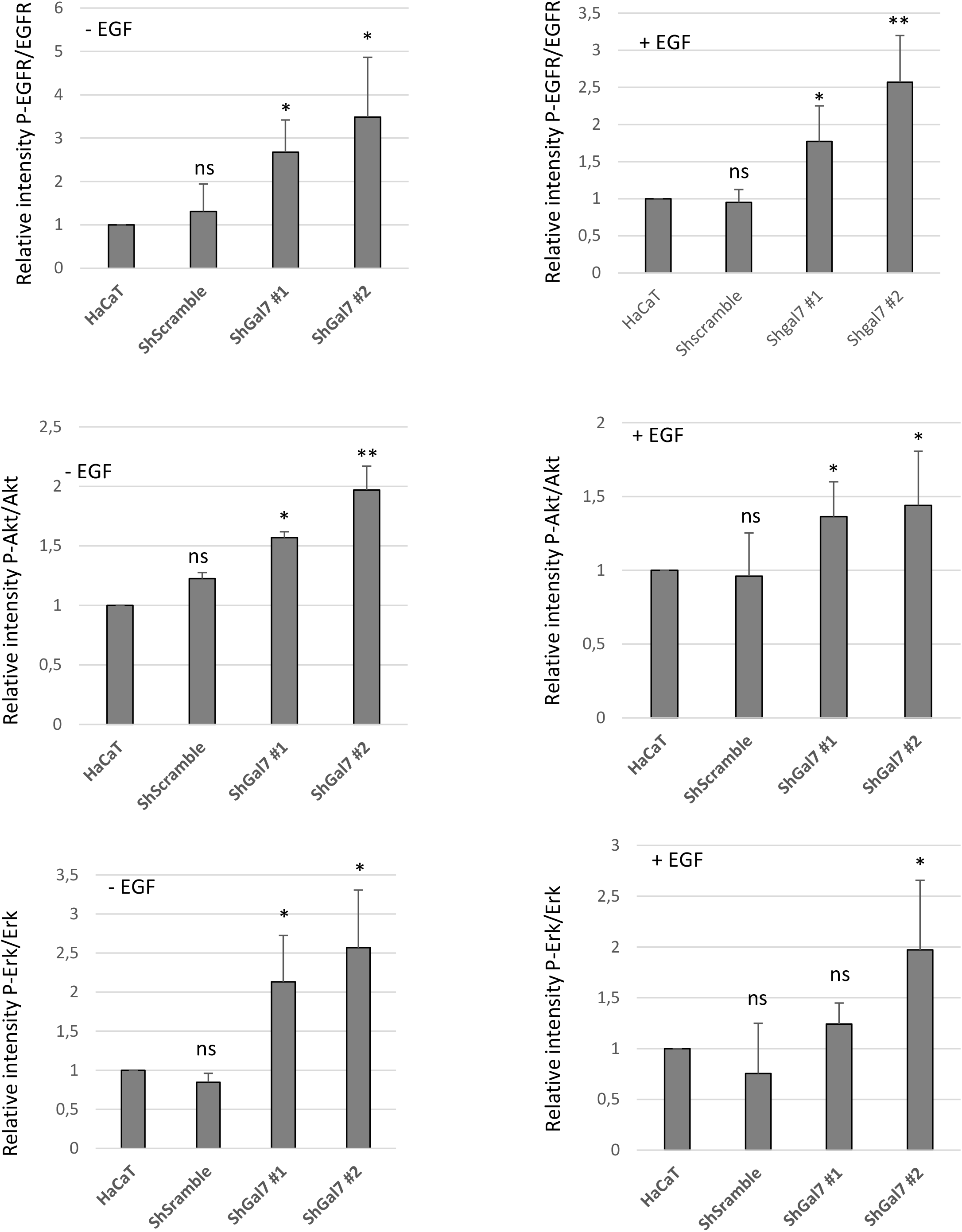
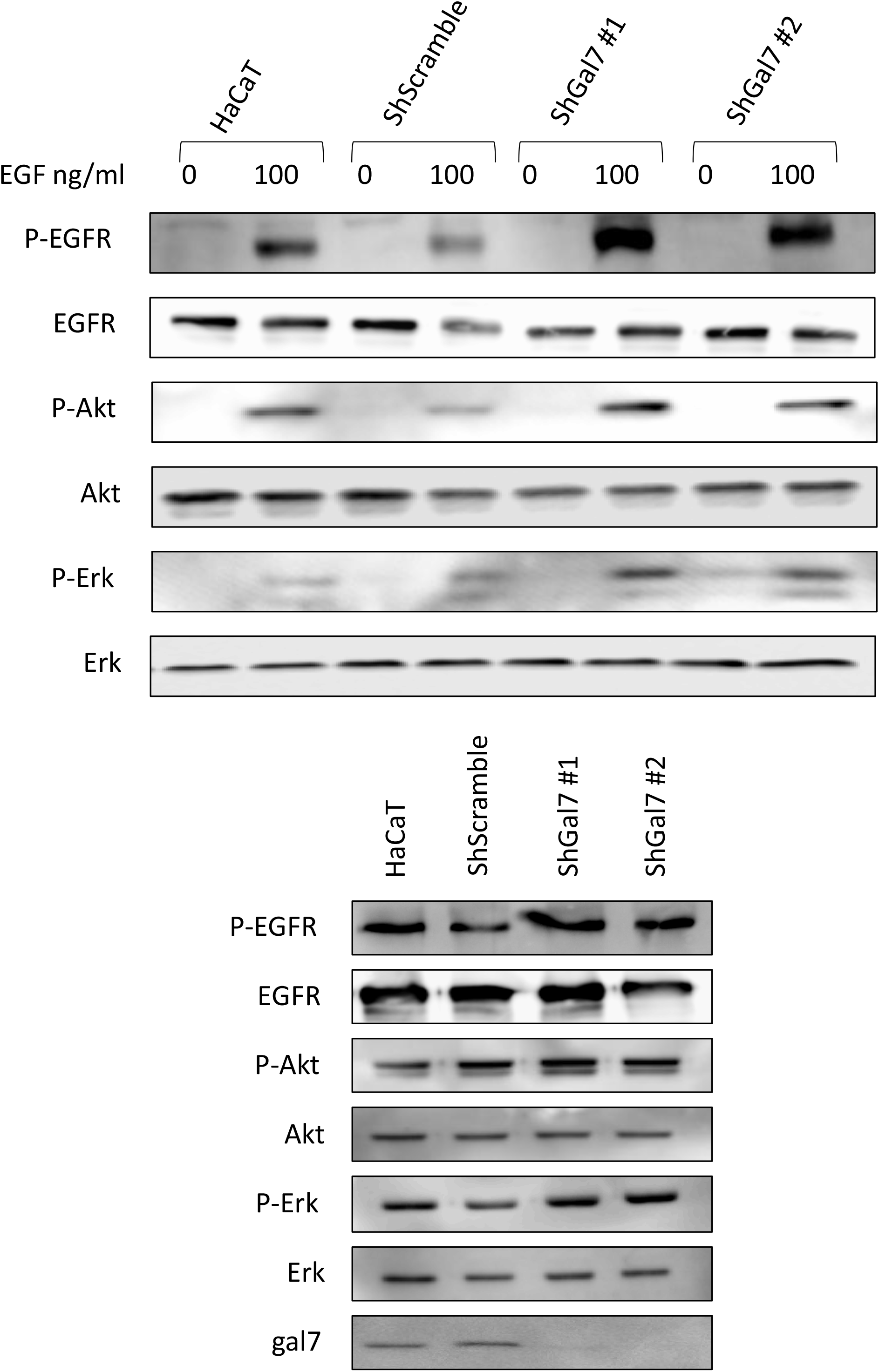
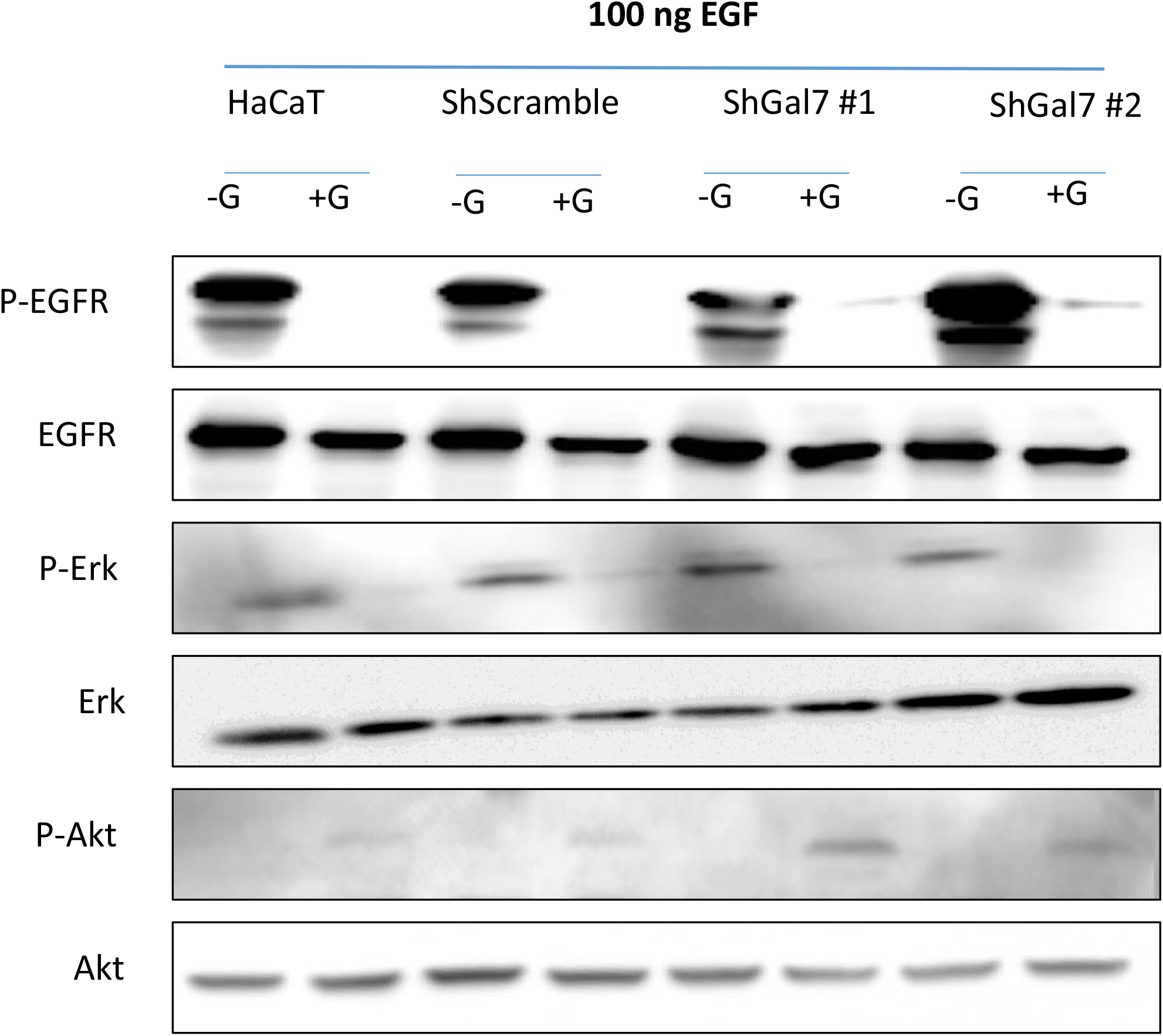
Aberrant EGFR phosphorylation by galectin-7 deficient cells is the cause for alteration of downstream pathways. A/ B/ Cells were treated or not with EGF (100ng/mL) for 15mn before being lysed. Immunoblots were probed for phospho EGFR (Y1068), total EGFR, phospho-ERK (p-ERK) total ERK, phopho-Akt and total Akt. C/ Cells were incubated with EGF and with 15μM of gefitinib for 15mn before being lysed. Quantification of the blots are representative of at least 3 independant expriments. * p<0,1 **p<0,05 ***p<0,001

When adding 100ng of EGF for 15min, a strong increase of EGFR phosphorylation was observed both in presence and in absence of galectin-7. For instance, the activation of phosphorylation of EGFR upon addition of EGF showed a 37 fold increase in HaCaT cells (figure S3). In absence of galectin-7 EGFR phosphorylation as well as Akt and Erk pathways showed the same tendency in the different conditions as in absence of EGF reinforcing the previous data showing that galectin-7 impairs EGFR phosphorylation and its downstream pathways. However these increases seem to be limited in presence of EGF which could be due to an activation plateau). In these experimental conditions we also detected an increase of STAT3 pathway (figure S2B) but Src pathway was still not activated (figure S2B In order to reinforce these observations we repeated the experiments in presence or in absence of gefitinib, a specific inhibitor of EGFR tyrosine kinase activity hence ensuring that indeed they do depend on EGFR activity. As shown in figures 3C and S2A, B, all the described pathways except Src are activated in a stronger manner in the absence of galectin-7. These results hence show that galectin-7 by interacting with EGFR restrains its constitutive and EGF-inductible phosphorylation and its downstream pathways.

As phosphorylation level has multiple consequences on receptors, we examined these different parameters.

### Galectin-7 depletion favors EGFR ubiquitination but not degradation

EGF treatment induces EGF receptor phosphorylation but also ubiquitination. Indeed signaling receptors are tightly regulated by this posttranslational modification, ubiquitination contributing to receptor endocytosis, sorting, and downregulation (Haglund and Dikic, 2012). Thus we investigated if galectin-7 could influence EGFR ubiquitination as a consequence of the modified phosphorylation pattern. We therefore performed immunoprecipitation experiments on cell lysates from HaCaT and ShGal-7 after EGF treatment during the indicated times, revealing that, as for phosphorylation (figure 4A), total EGFR ubiquitination was more intense in cells deprived for galectin-7 especially after 5min of stimulation (figure 4A). This led us to suppose that galectin-7 would negatively regulate EGFR ubiquitination. In fact ubiquitination after ligand binding plays a fundamental role both in EGFR endocytosis and intracellular sorting. Indeed, EGFR ubiquitination starts at the plasma membrane and continues along the endocytic pathway. Ubiquitination is also critical at later steps targeting EGFR for degradation through trafficking to lysosomes (Conte and Sigismund, 2016). Thus, we wondered if galectin-7 influences EGFR degradation.

**Figure 4:**
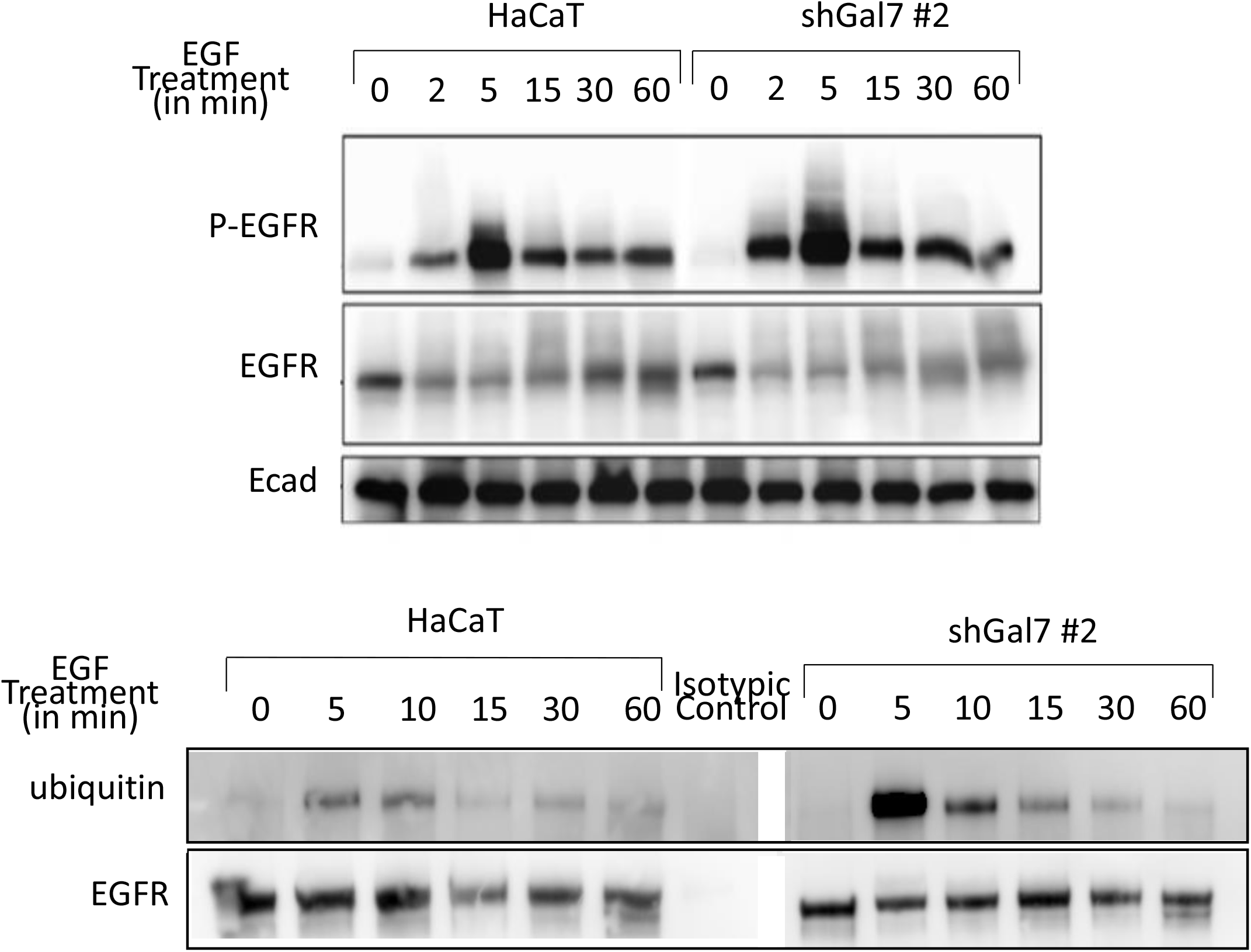
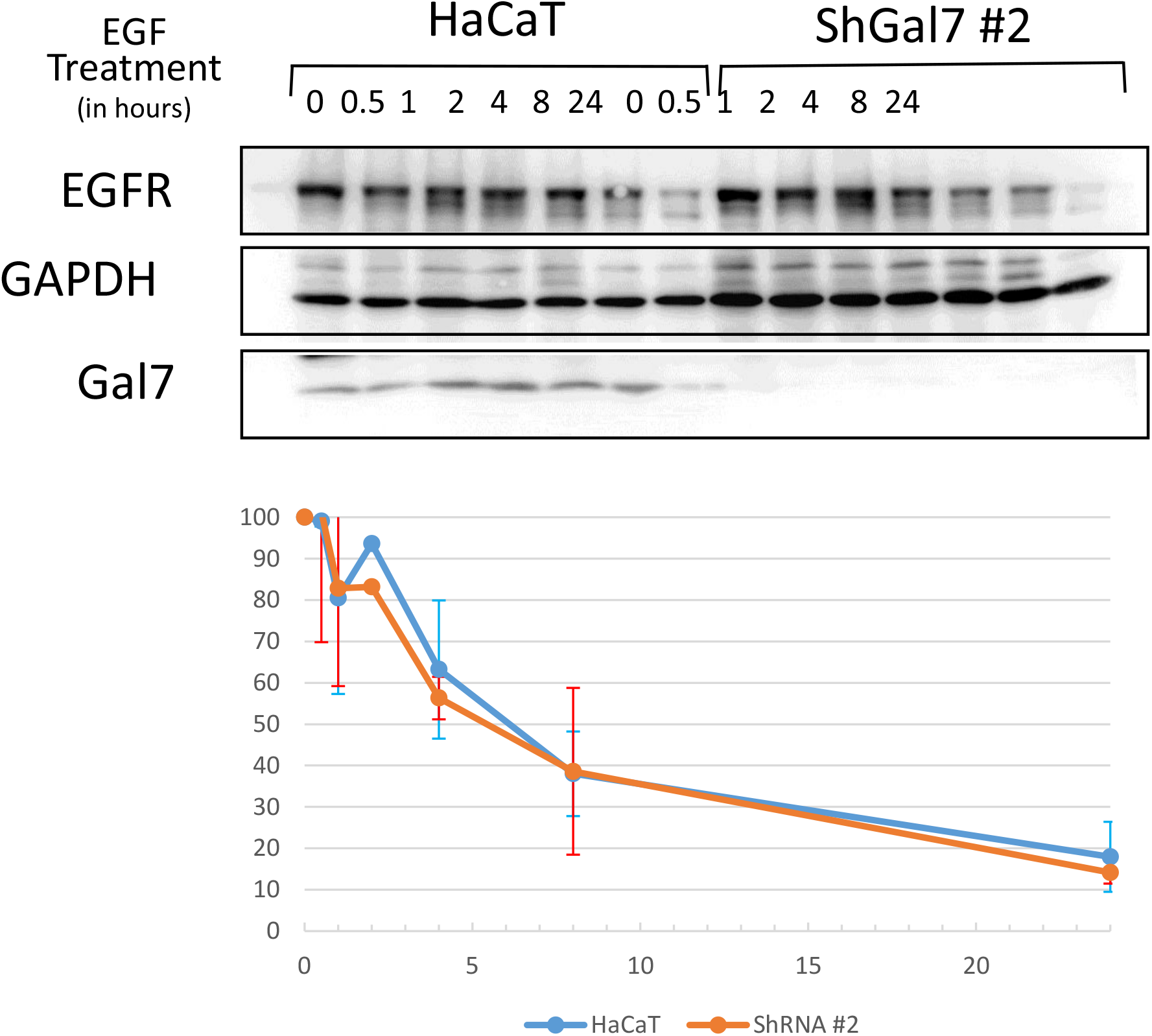
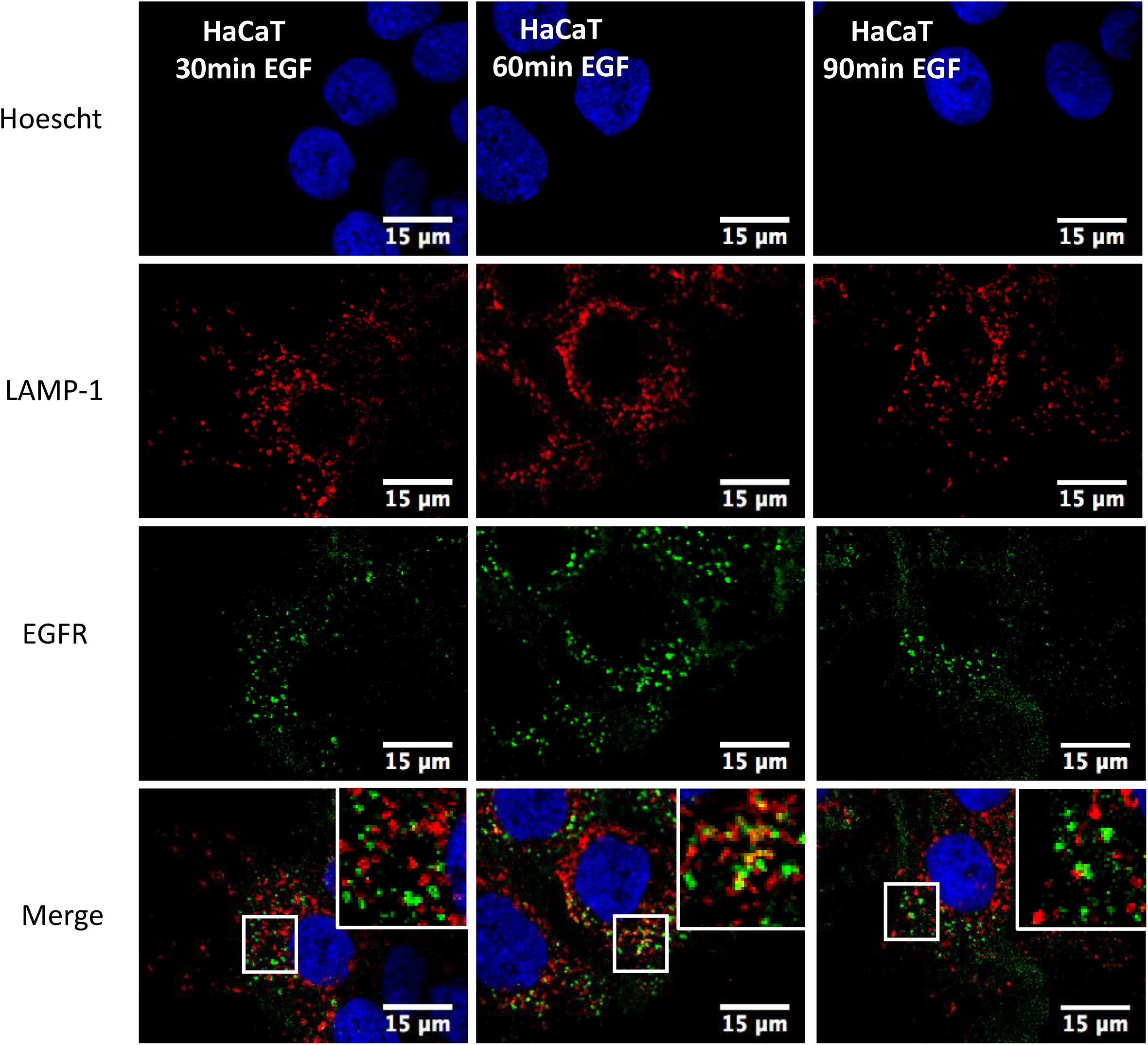
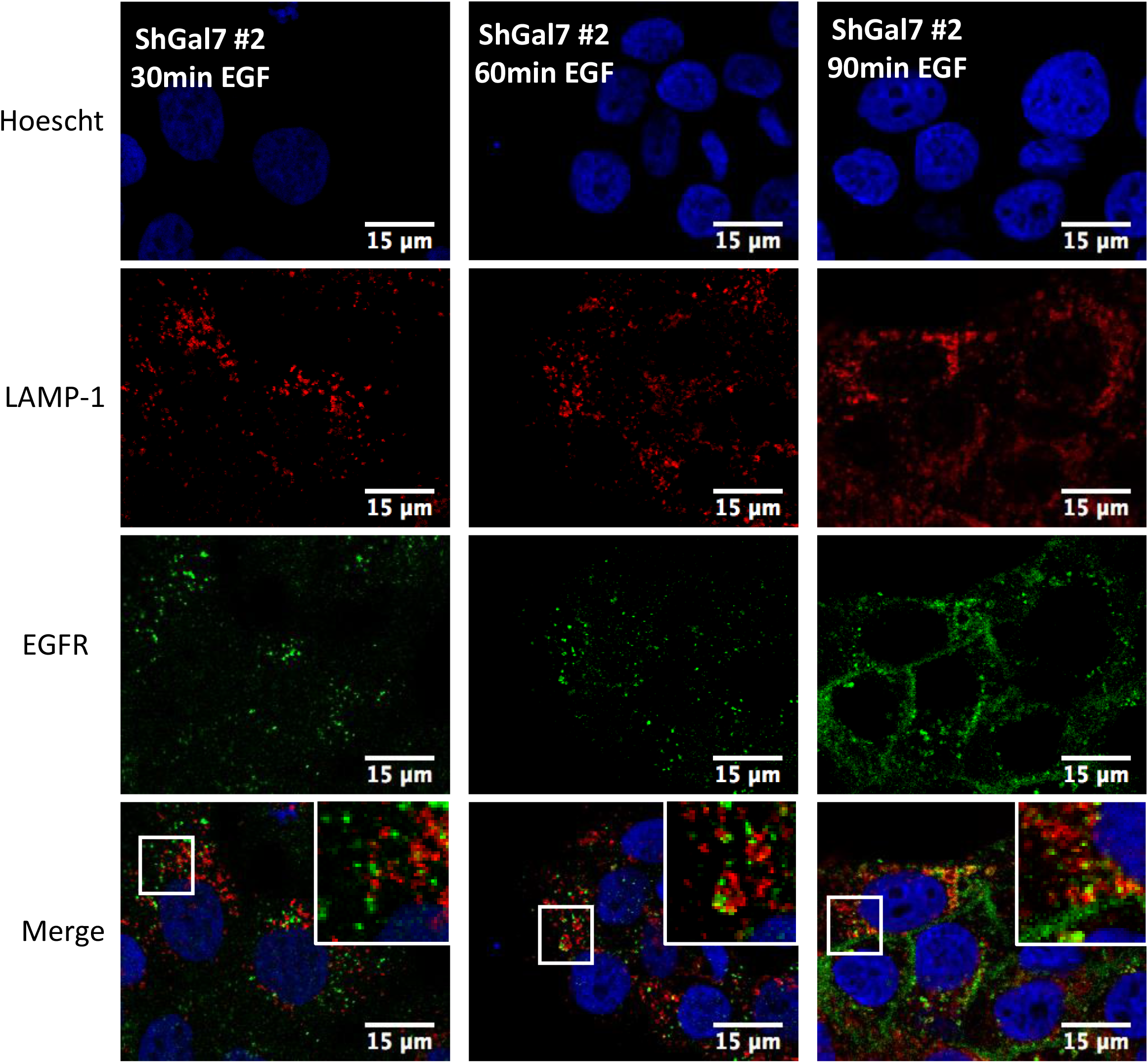
Upregulation of EGFR phosphorylation and ubiquitination in absence of galectin-7 after EGF treatment. A/ Immunoprecipitation of total EGFR from whole cell extracts of HaCaT or shGal-7 cells obtained after treatment with 100ng/ml EGF for the indicated times. Immunoblots were probed for ubiquitine and total EGFR. Time course of EGFR activation by EGF in HaCat or shGal-7 cells treated with 100 ng/ml EGF. Whole cell extracts were prepared at the indicated times and analyzed by immunoblotting for P-EGFR and total EGFR B/EGFR Protein Stability after EGF treatment. For EGFR stability experiments, HaCaT cells were plated in 6wells plates, starved overnight then treated with EGF 100ng/ml with or without Cycloheximide at 25μg/ml. After the indicated times, cells were lysed and total EGFR levels were detected by immunoblotting. C/ Representative immunofluorescence of EGFR (green) and LAMP-1 (red) in HaCaT cells and in ShGal7 #2 after 30, 60 and 90 min of treatment with 100ng/mL of EGF. Nucleus are stained in blue (Hoescht) Scale bar=15μM

To determine EGFR stability, we checked the level of EGFR present in cells at different times of EGF treatment in presence of cycloheximide. Cycloheximide inhibits translation thus the EGFR neo synthesis that could otherwise mask degradation. However we observed no significant differences in EGFR degradation in our assays between cells depleted or not in galectin-7 (figure 4B). These results have been confirmed by immunofluorescence through co-staining of EGFR and LAMP-1 at 30mn, 1hr or 1hr30min (figure 4C). Thus, it can be seen that in both cases EGFR mainly seems to be co-localized with lamp-1 after 1 hour of stimulation with EGF. Moreover, in absence of galectin-7 EGFR is not massively targeted to lysosomal compartments. Thus galectin7 would not would not be implicated in EGFR degradation.

### Galectin-7 impacts EGFR trafficking

The previous results led us to hypothesize that ubiquitination modification would influence the endocytosis of EGFR in cells. In order to decipher the consequences of galectin-7 depletion we studied EGFR intracellular trafficking by co-staining with several intracellular compartments. To this purpose, pulse-chase experiments were conducted stimulating cells for 20 min with 100 ng ml^−1^ conjugated-EGF (or galectine-7) and after removing unbound ligand chased for 5min to 4 hours. Immunofluorescence staining followed by colocalization analysis establishes the endocytic trafficking of EGF ligand through the degradative pathway from early (TfR positive compartments) to late (CD63 positive compartments) within 2 hours. At late stages (60-90min) EGFR partially colocalizes with CD63 in both HaCaT cell line and ShGal7 cells indicating that EGFR is indeed able to reach the late endosomal compartments (figure 5A). As ubiquitination is also implicated in the first steps of endocytosis and because EGFR degradation is not impacted in cells depleted of galectin-7, we investigated if EGFR could be more efficiently recycled to the plasma membrane. We incubated the cells with fluorescent transferrin which uptake allowed us to explore early and recycling endosomes. Hence more colocalization is detected in cells lacking galectin-7 suggesting that a high amount of the endocytosed P-EGFR in mutant cells is recycled back to the plasma membrane (figure 5B). Thus during EGF stimulation, galectin-7 restrains P-EGFR endocytosis and its recycling at the plasma membrane.

**Figure 5:**
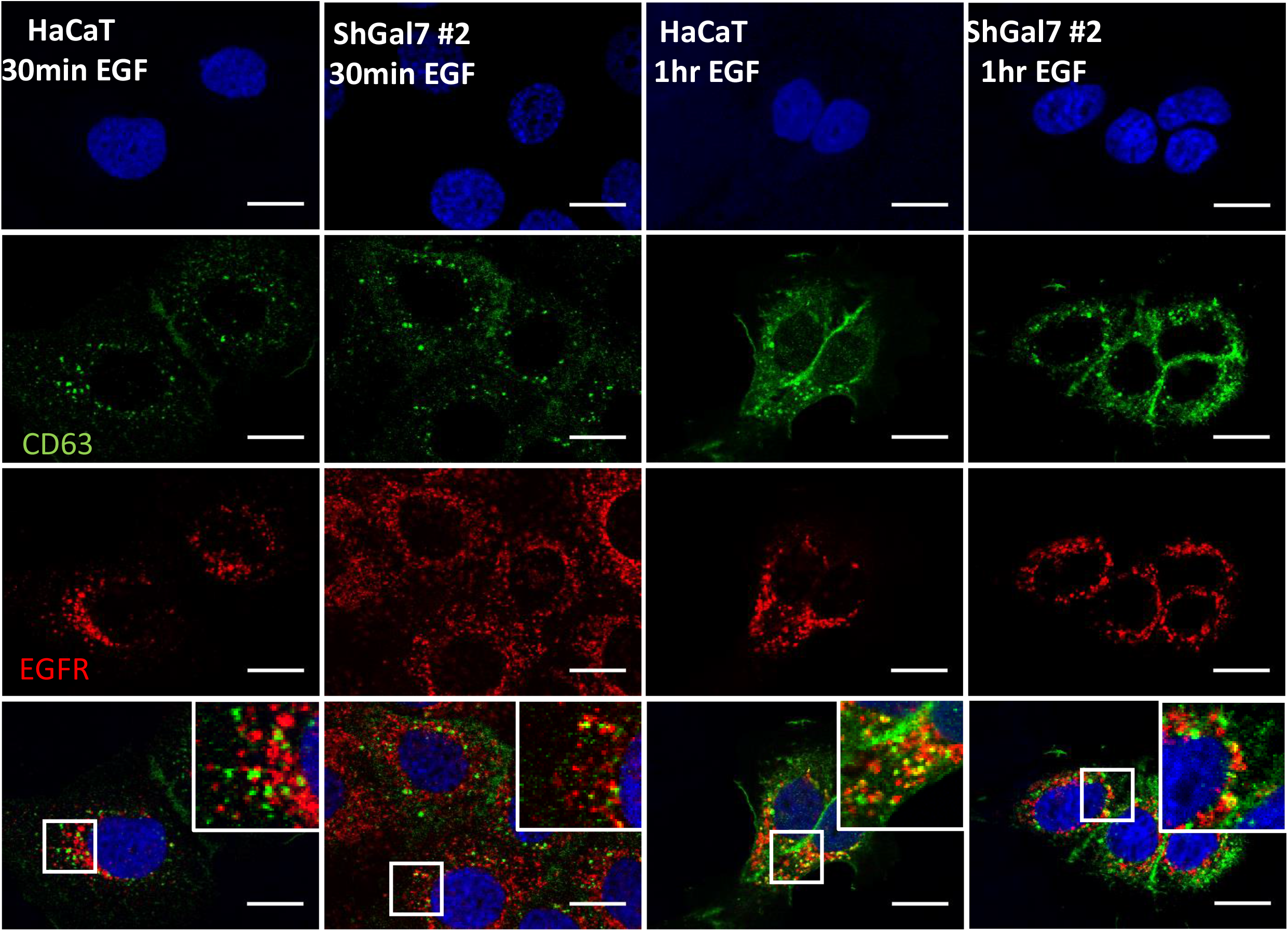
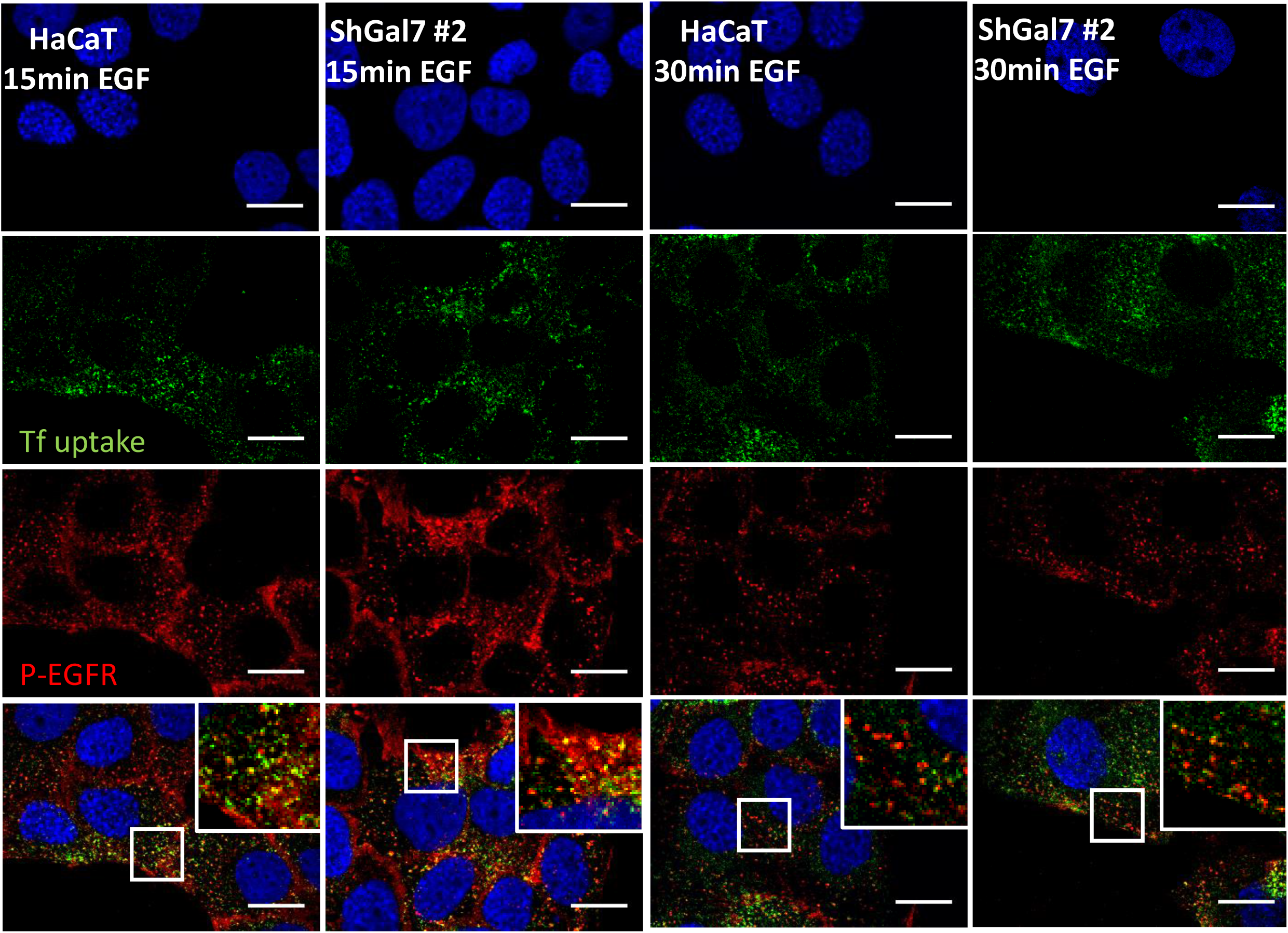
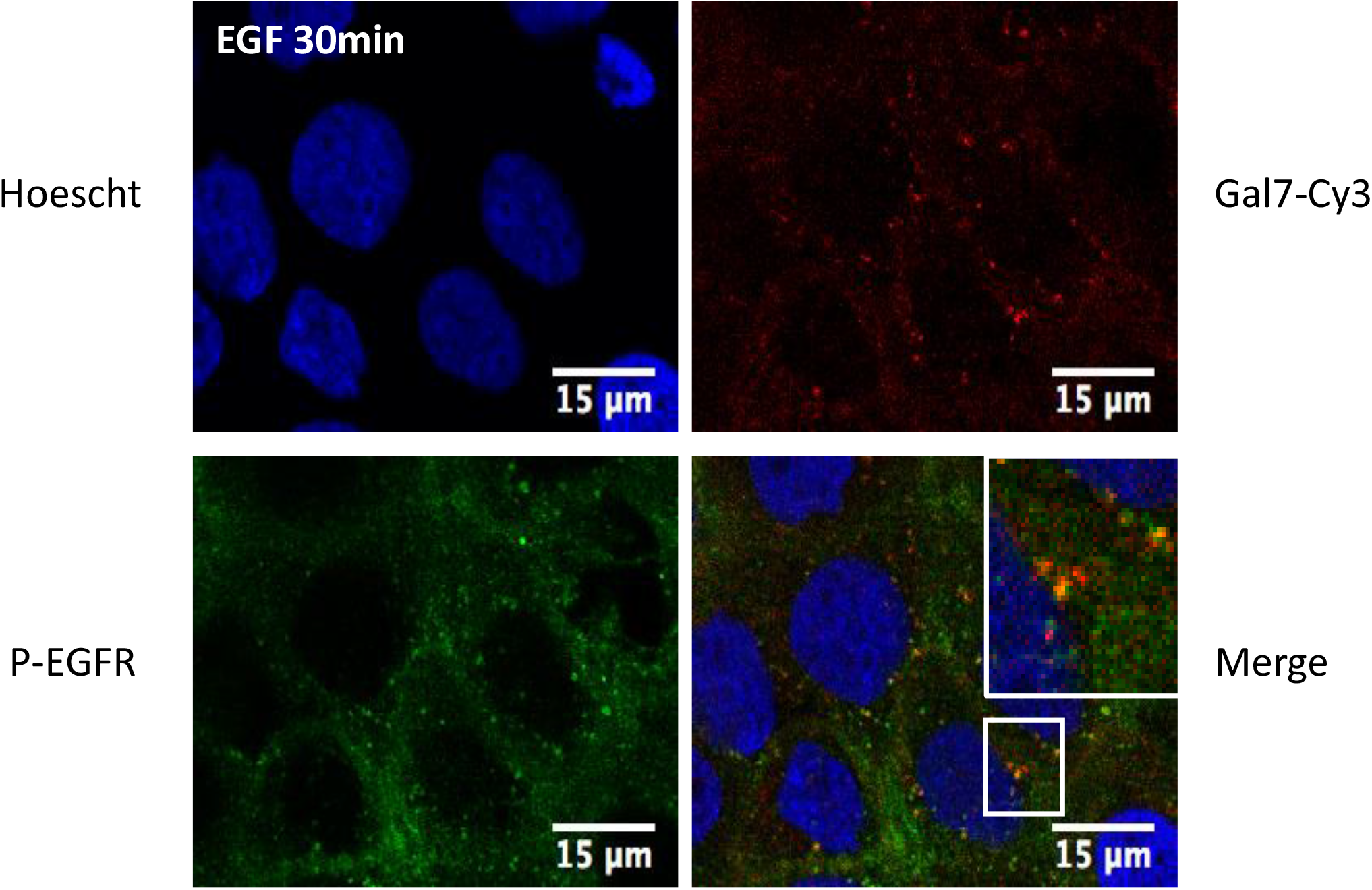
Galectin-7 is endocytosed with EGFR and modulates EGFR endocytosis. A/ Representative immunofluorescence of CD63 (green) and EGFR (red) after 30min and 1hr of treatment with EGF Nucleus are stained in blue (Hoescht) Scale bar=15μM B/ Representative immunofluorescence of transferrin (green) and P-EGFR (red) in HaCaT cells and in ShGal7 #2 cells at 30, 60 and 90min after treatment with 100ng/ml of EGF. Nucleus are stained in blue (Hoescht) Scale bar=15μM C/ Representative immunofluorescence of galectin-7 (red) and P-EGFR (green) in HaCaT cells after 30mn of EGF treatment at 100ng/ml. Nucleus are stained in blue (Hoescht) Scale bar=15μM.

Interestingly, as described above in proximity ligation assay, we observed an interaction between galectin-7 and EGFR in cell cytoplasm. To confirm these results we used recombinant galectin-7 coupled to Cy3 (rgal7-Cy3) and performed co-staining with P-EGFR. Strikingly, recombinant galectin-7 was repeatedly observed in endocytosed P-EGFR-containing vesicles (Figure 5C), indicating that galectin-7 was probably co-endocytosed with P-EGFR. On the contrary, colocalization assays of gal7 and LAMP-1 gave no results (data not shown), letting us consider that galectin-7 could travel with P-EGFR during the early steps of endocytosis. All these results made us consider that galectine-7 exerts a negative control on EGFR endocytosis and recycling.

### Galectin-7 impacts cell migration and cell proliferation in absence of EGF

As galectin-7 restrains EGFR phosphorylation and signaling pathway even in absence of added EGF we wondered if galectin-7 could also alter cell motility and cell proliferation. So we set up wound healing assay using insert removal technics (Advedissian et al., 2017b) to investigate the migratory potential of galectin-7 depleted clones in cell starvation conditions. Sixteen hours after insert removal, the ShGal7 clones exhibited a significant increase in wound healing capacity of respectively 60% (ShGal7 #1) and 90% (ShGal7 #2) compared to control HaCaT cells (Figure 6A, 6B). Interestingly when adding soluble recombinant galectin-7 to ShGal7 clones we can observe that cell migration is similar to control cells (figure 6A). Hence this rescue reinforces the hypothesis that galectin-7 also exercises a negative control over cell migration capacity in these conditions.

**Figure 6:**
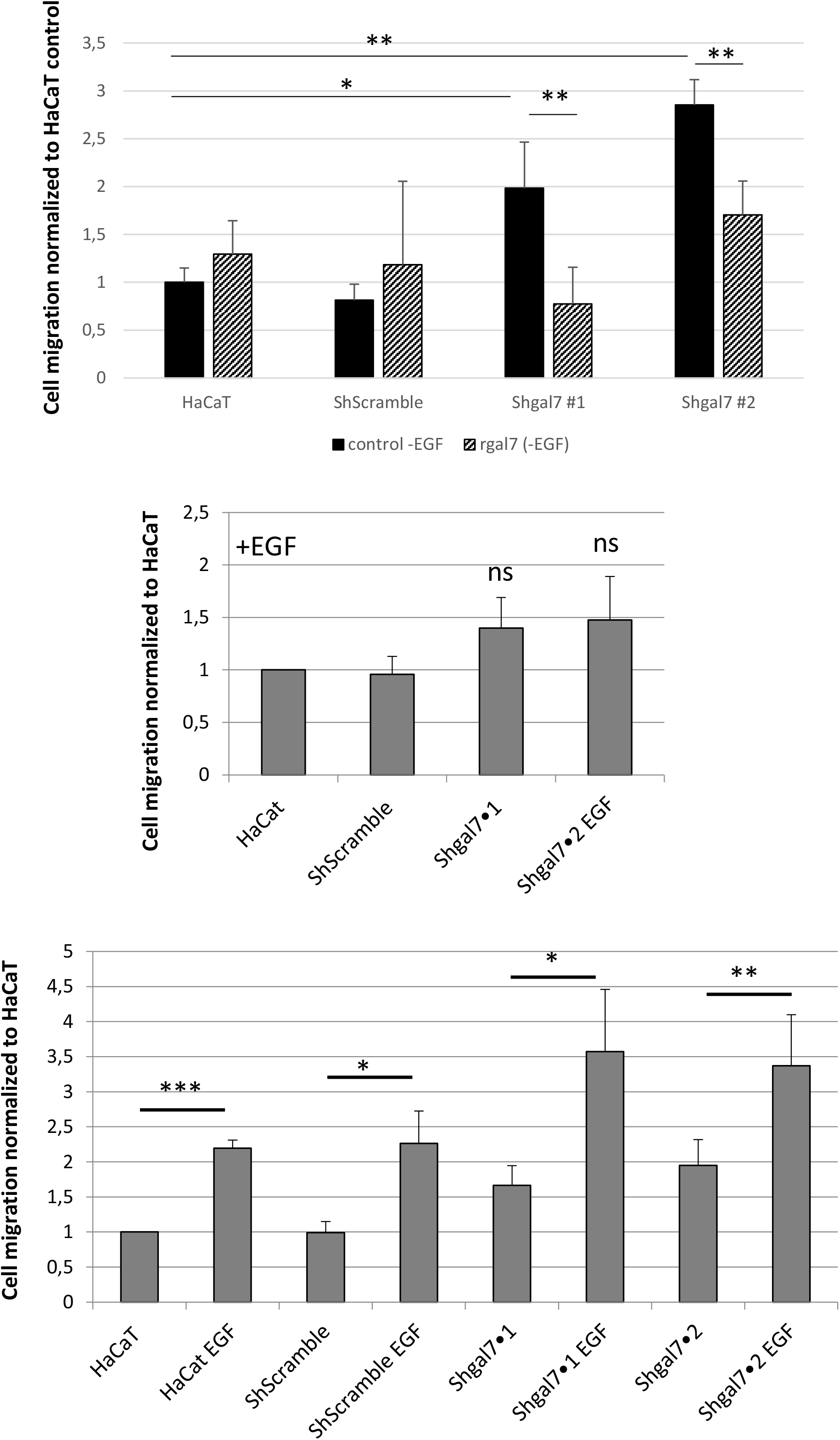
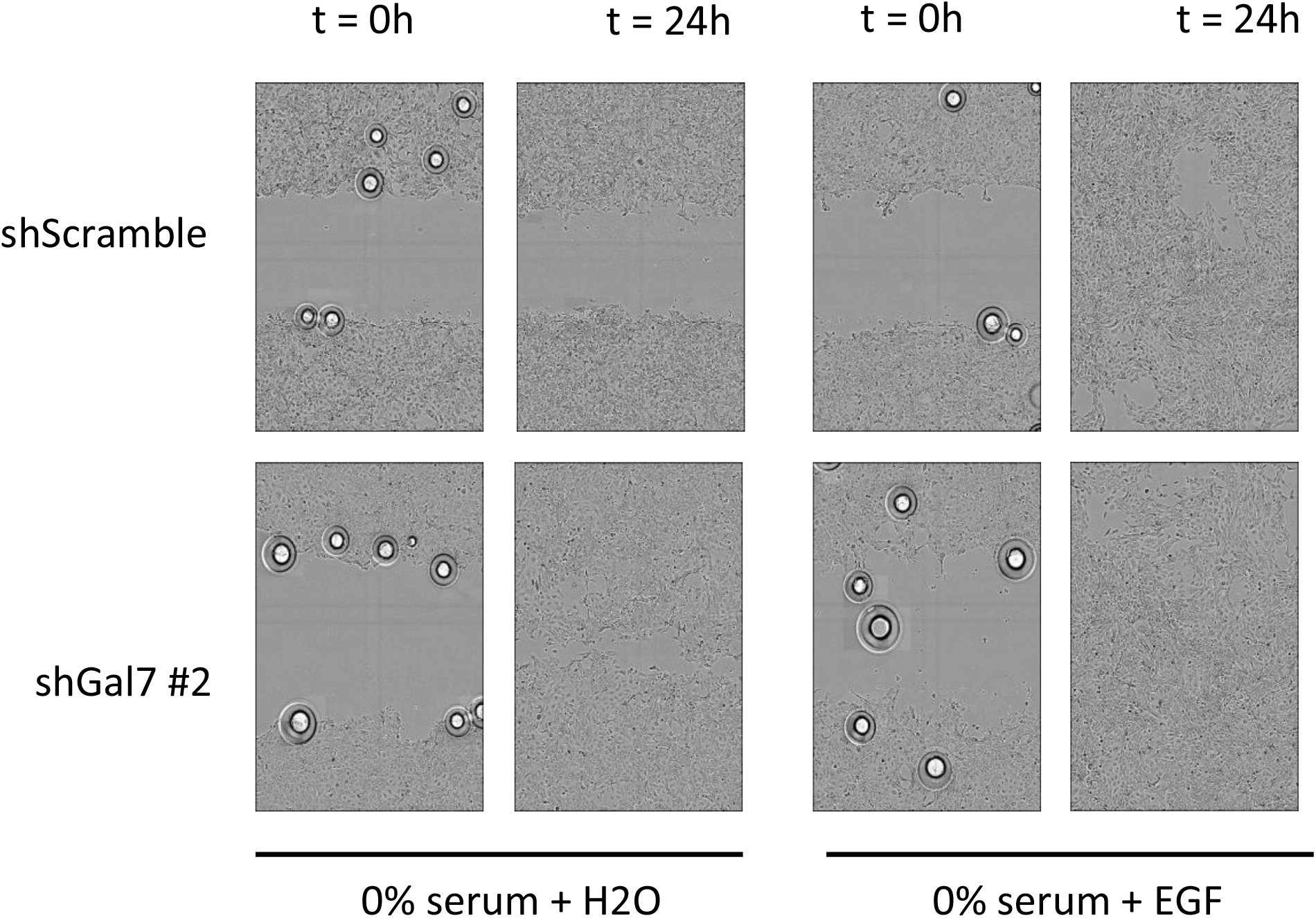
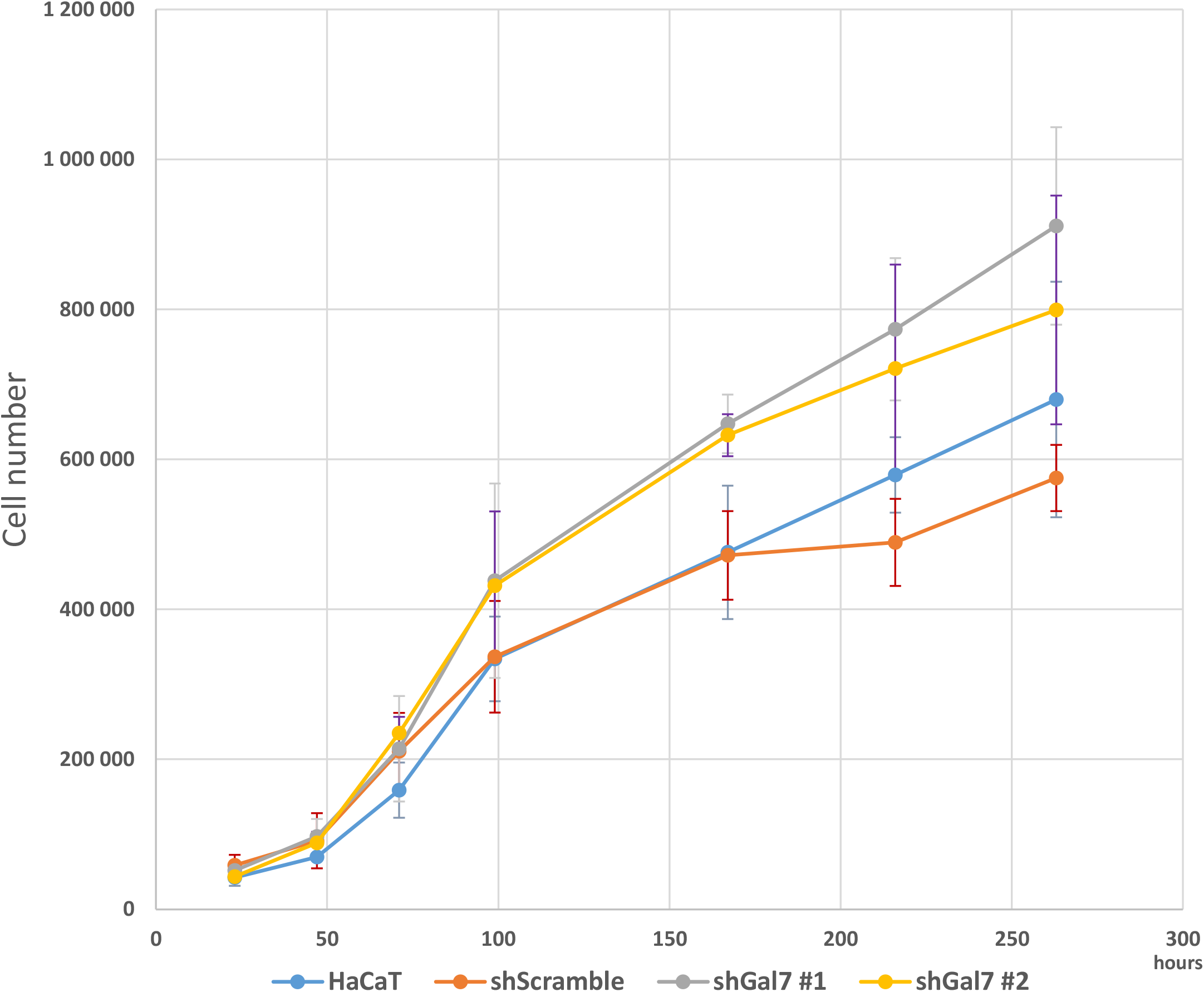
Galectin-7 downregulation enhances cell migration and proliferation of HaCaT keratinocytes. A/ Percentage of wound closure normalized to HaCaT WT cells in insert removal wound healing experiments. Rescue experiments have been conducted with 0.5mM of recombinant galectin-7. Mean ± s.e.m. are represented (n=4). B/ Images have been extracted from videos of cell migration. Magnification 20x. C/ Curves are the results of total cell proliferation assays of HaCaT and shGal7 #2 cell lines counted during eleven days to calculate the mean ± standard deviation (SD). Results are mean of three independent experiments performed in duplicate.

**Figure 7:**
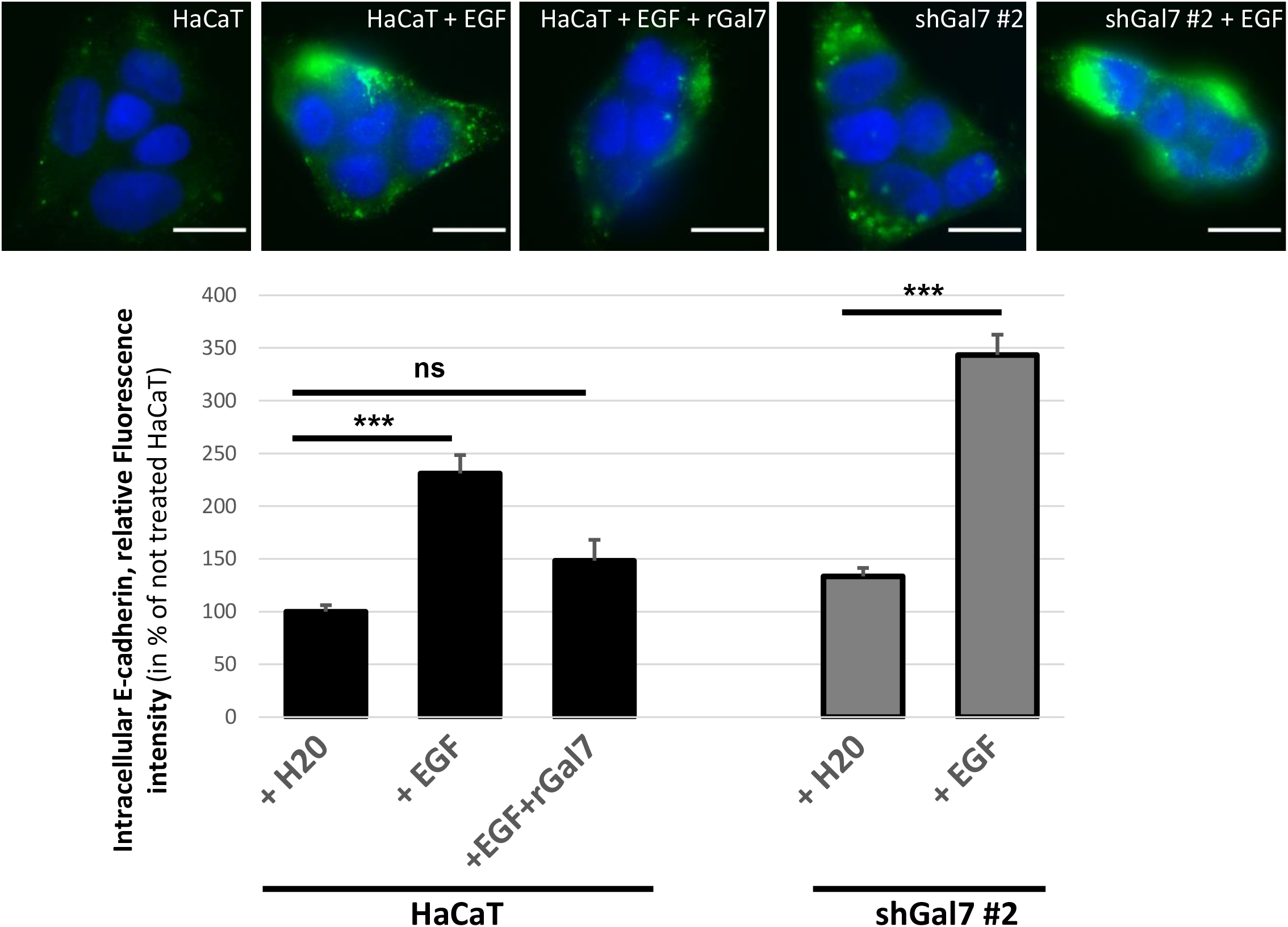
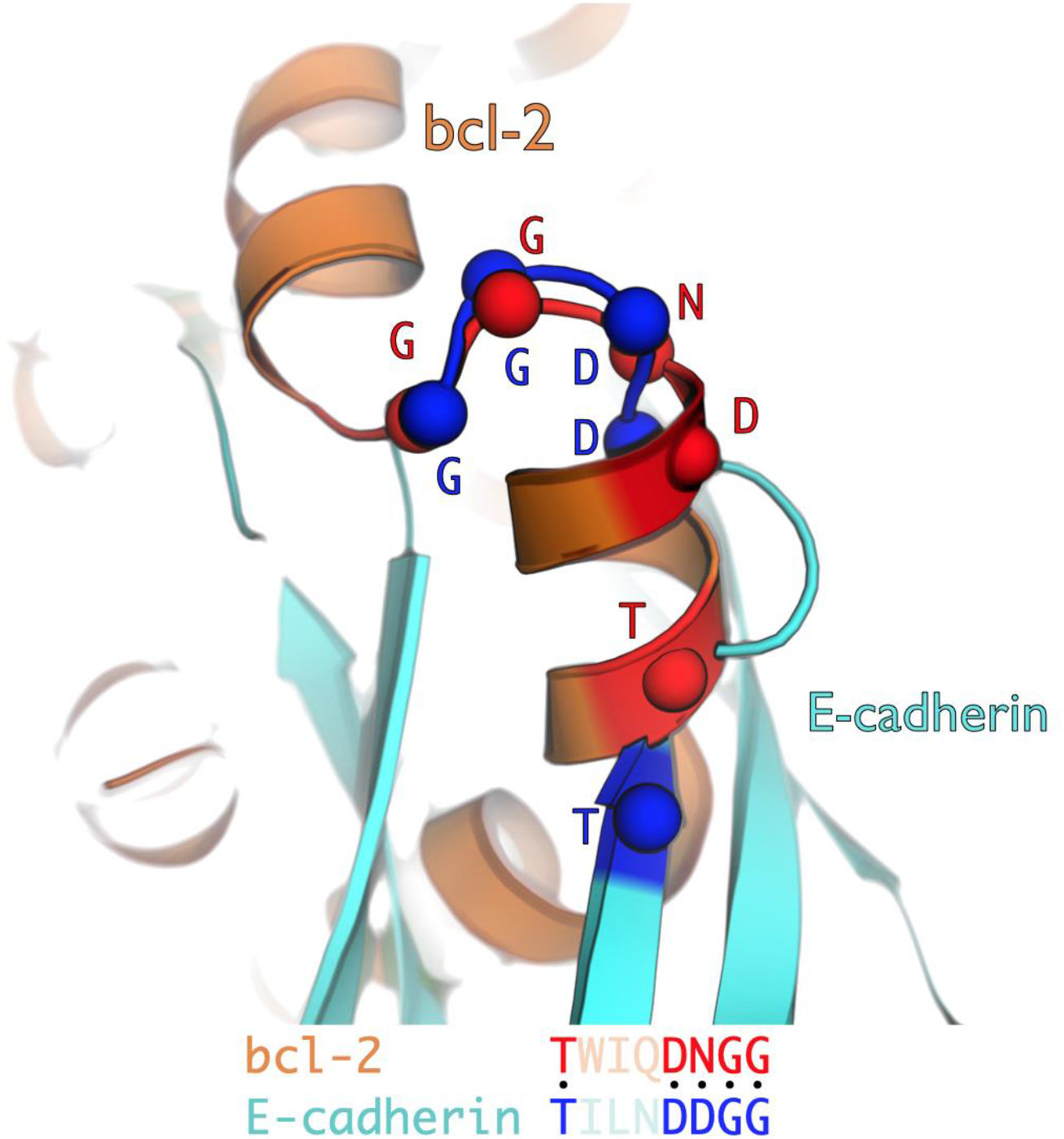
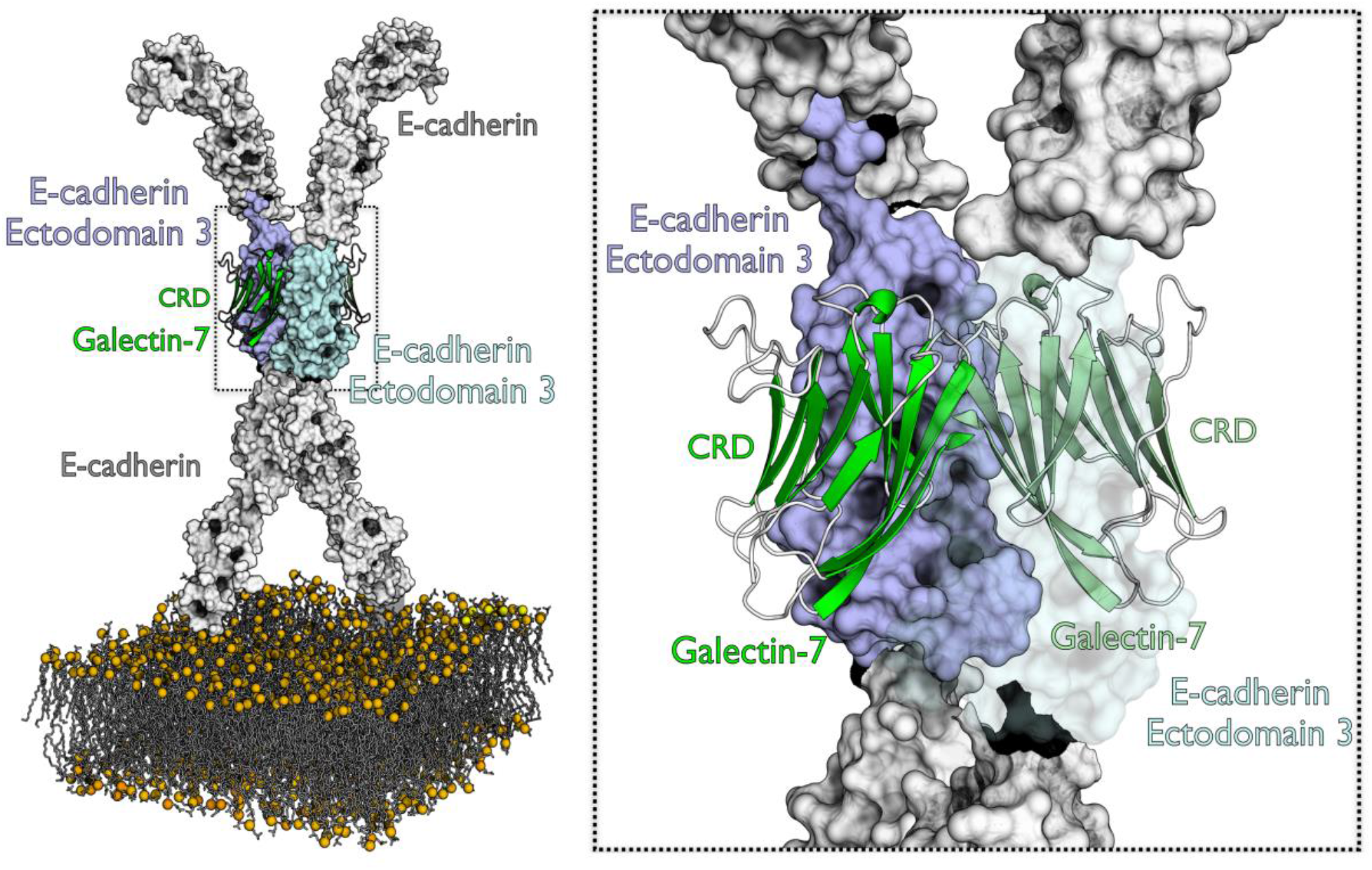
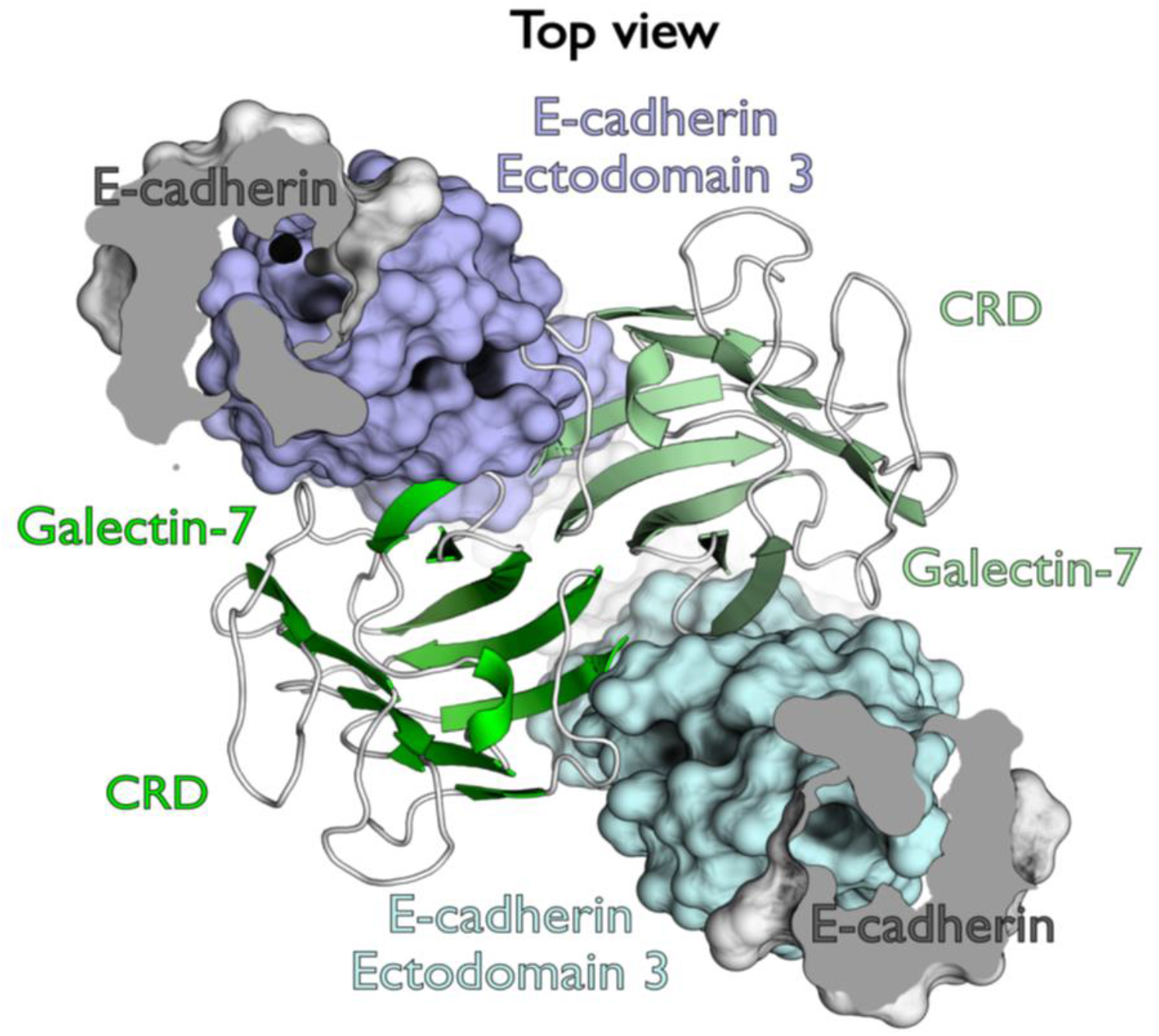
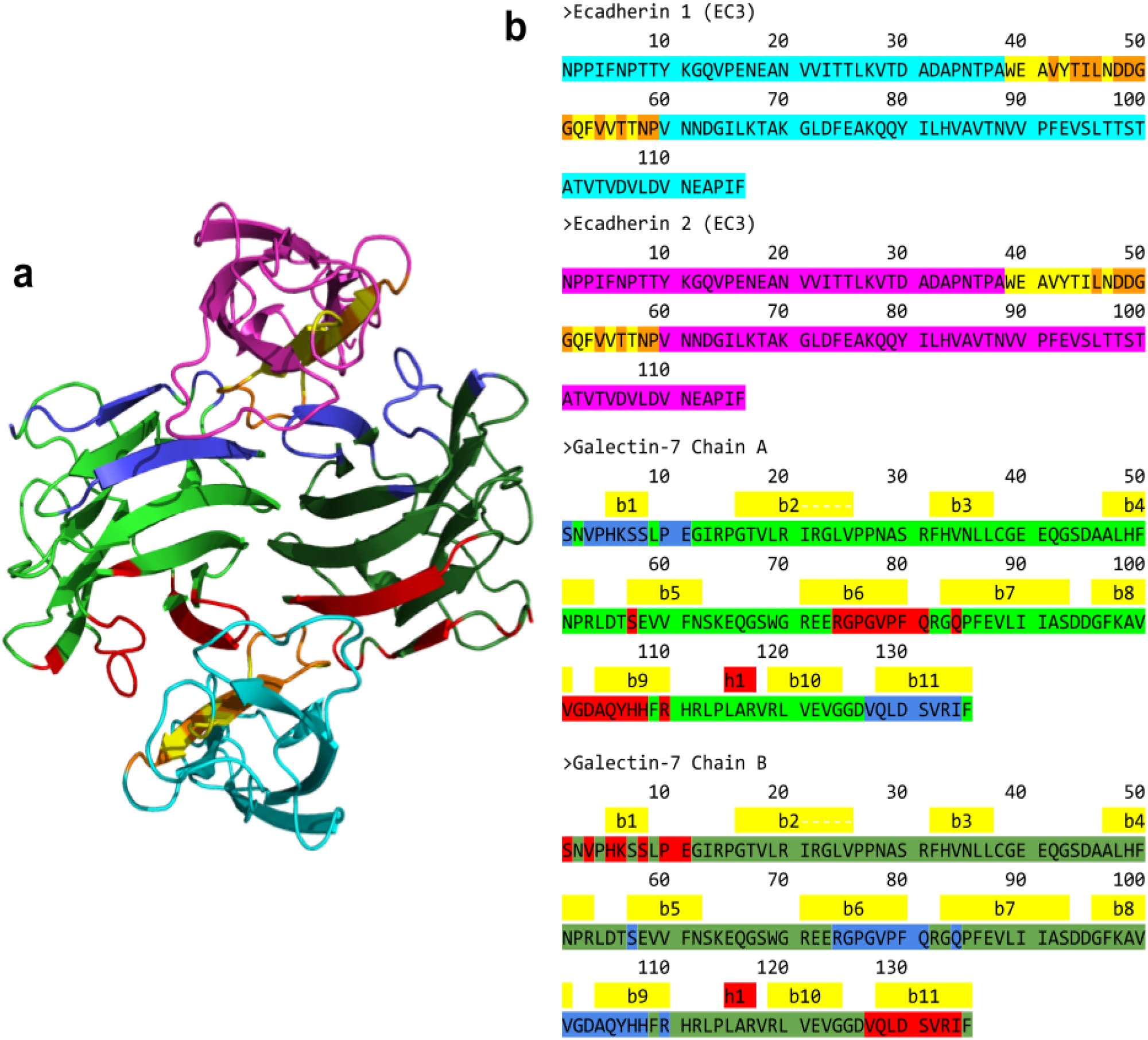
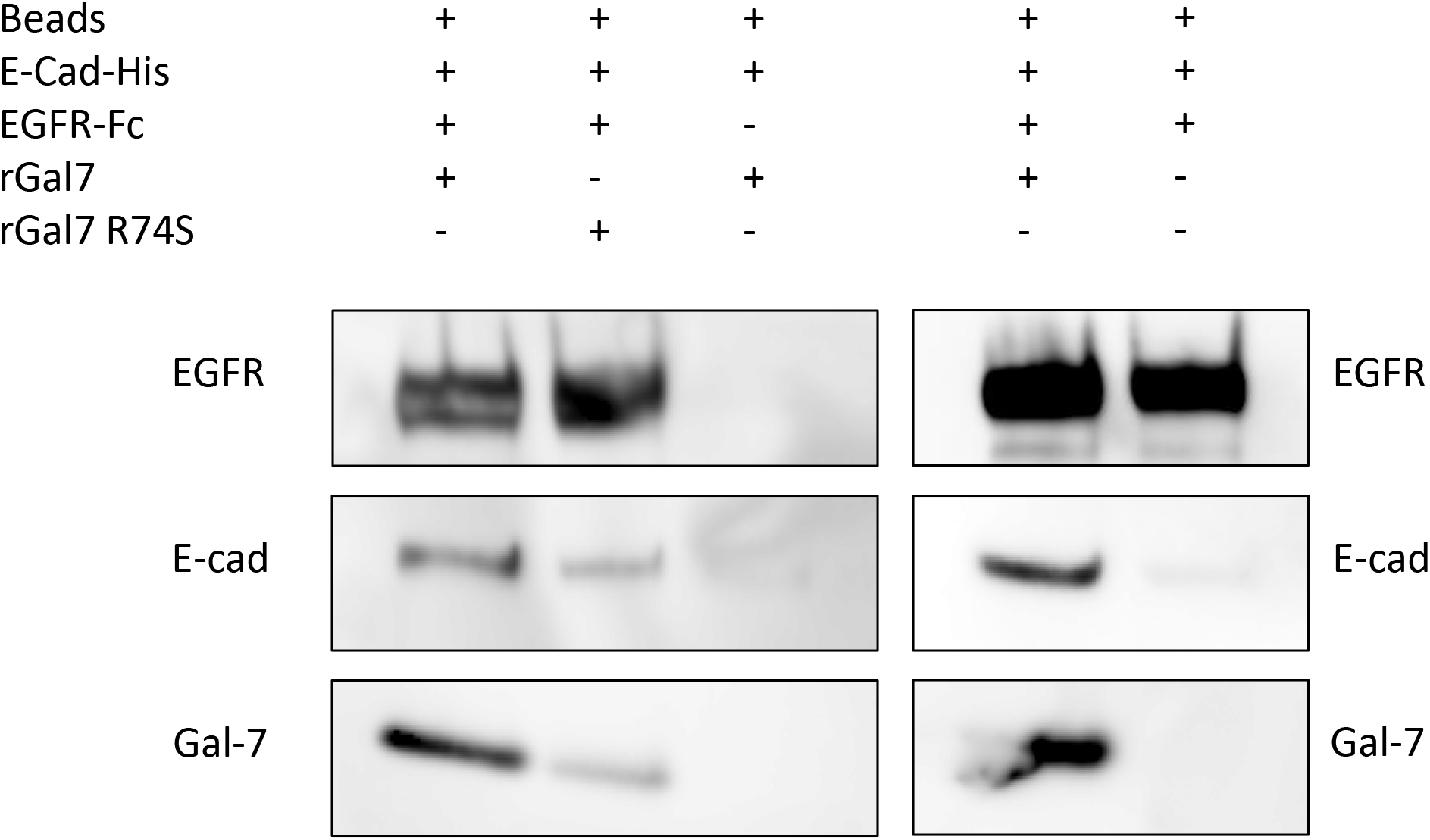
Galectin-7 interacts with both EGFR and E-cadherin. A/ Modulation of E-cadherin internalization by galectin-7 and EGF. Representative images of HaCaT and ShGal7 #2 after E-cadherin antibody uptake for 30 min. Cells were previously treated or not with 100ng/mL of EGF. Histograms represent corresponding quantifications in percentage reported to HaCaT not treated cells. B/ Structural alignment of sequence motifs found in E-cadherin ectodomain 3 and Bcl-2. The threonine residues, as well as the DXGG motif can be structurally aligned with high precision. Ca of aligned residues are represented as spheres. C/ *In silico* model of interaction of two E-cadherin molecules on both sides of galectin-7 homodimer placed on the plasma membrane. Galectin-7 chains are presented in green shades, E-cadherin ectodomains in light grey. Ectodomains 3 are colored in pale violet and blue. A zoom on the binding region can be seen on the right. On the zoom, pale blue E-cadherin 3 ectodomain is displayed in transparency mode to better observe the interaction interface. D/ Top view of two E-cadherin molecules interacting with galectin-7. The same color scheme of figure 2 have been used. The two N-ter E-cadherin ectodomains are cut for clarity. E/ a) Firsts docking poses on the galectin-7 dimer for the first and second ectodomain 3 (EC3) of e-cadherin, in cyan and magenta, respectively. Chains of galectin-7 dimer are represented in green shades. Sequence motif identified in e-cadherin in colored in yellow. In e-cadherin, interface frequent residues are colored in orange. Interface frequent residues made by galectin-7 with the first e-cadherin molecule are in red; those with the second molecule are in blue. b) Sequences of the two EC3 molecules and the galectin-7 dimer are displayed. Sequences and contacts are colored as described above. F/ *In vitro* binding assays were performed using recombinant wild-type human galectin-7 (rGal7); CRD mutated human galectin-7 (R74S), Extracellular domain of human E-cadherin fused to a His tag (E-cad-his) and extracellular domain of human EGFR fused to human IgG1 Fc fragment (EGFR-Fc). E-cadherin precipitated with EGFR-Fc only in presence of galectin-7.

Similarly, we performed cell proliferation assay seeding the different cell lines at the same confluence and evaluated cell population size regularly during more than 10 days. A statistical difference is already detectable after one week in culture with a higher number of cells counted in both ShGal7 clones compared to the HaCaT cell lines. We can observe on figure 6C that HaCaT cells need about an additional 48hrs to multiply their population by 3. Thus galectin-7 inhibits growth factor independent cell proliferation. These observations are consistent with the above reported role of galectin-7 reducing the basal activation level of the EGFR pathway.

### Galectin-7 connects EGFR and E-cadherin and interfere with EGF dependent E-cadherin dynamics

We previously documented that galectin-7 regulates E-cadherin dynamics at the plasma membrane (Advedissian et al., 2017b). EGFR and E-cadherin are known to interact together with consequences on the balance between proliferation and differentiation as illustrated by the results of Wilding et al., (Wilding et al., 1996). The reported results indicate that overexpression of the EGFR correlates with perturbation of the E-cadherin/catenin complex suggesting an underlying functional interaction between growth-regulatory factors and E-cadherin. Hence, we decided to study a potential bridge between the three molecules. We first ascertain that E-cadherin and EGFR co-precipitate in HaCaT Cells (Figure S4). We then performed internalization assay of E-cadherin under the dependence of EGF. We measured the internal intensity of fluorescence corresponding to E-cadherin using an anti-E-cadherin extracellular domain HECD-1 antibody. As previously described, at basal level, E-cadherin is internalized with better efficiency in absence of galectin-7. In presence of 100ng/ml EGF we observed that E-cadherin is 2.3 times more internalized in HaCaT cells when compared to cells without EGF (figure 8A). Interestingly, addition of extracellular galectin-7 on HaCaT cells reduces the EGF induced endocytosis of E-cadherin rendering E-cadherin more resistant to EGF treatment hence confirming that galectin-7 impairs E-cadherin internalization (Advedissian et al., 2017b). In contrast, E-cadherin endocytosis is more intense in cells depleted of galectin-7. Thus galectin-7 has an effect antagonist of EGF on E-cadherin endocytosis and does negatively regulate the endocytosis of E cadherin with and without EGF. These results prompted us to consider that galectin-7 could form a complex with E-cadherin and EGFR thus co-regulating both these membrane proteins.

### *In silico* assessment and modeling of Galectin-7, E-cadherin complex and insights for EGFR interaction

In light of our findings, the binding between EGFR and galectin-7 seems dependent of the CRD suggesting that galectin-7 would directly interact with both EGFR and E-cadherin making a physical link between them. To further decipher and assess this interaction, we have conducted *in silico* modeling between E-cadherin and galectin-7 based on previous experiments (Advedissian et al., 2017b).

Because galectin-7 interacts with EGFR through its CRD domain, interaction between galectin-7 and E-cadherin should take place at distance of the CRD domain. This glycosylation-independent interaction between E-cadherin and galectin-7 thus might echo the Bcl-2/galectin-7 interaction. Interestingly, even if Bcl-2 and E-cadherin do not share global similarity, they share a common motif of 47 residues with 26% identity at the level of E-cadherin extracellular domain 4 which was identified by local alignment on E-cadherin extracellular sequence. Similarly, when aligning E-cadherin domain 3 with Bcl-2 using the same protocol, another motif of 25 residues can be identified which shares 30% identity between each other (Supp. figure. S5A). These results suggest that galectin-7 interaction with E-cadherin might be mediated by the latter domains 3 or 4. Docking simulations with either domain 3 or domain 4 allow to identify most probable interface residues. While recurring residues of domain 4 do not constitute a patch on the motif found by local sequence alignment, 50% of domain 3 motif residues are part of the interface in at least 1,000 docking poses. These residues, when mutated *in silico*, affect E-cadherin binding mode (data not shown), confirming their fundamental role in the interaction. Interestingly, the most impacting mutations are those affecting several residues at the same time, particularly those impacting the DDGG motif found in Bcl-2 (figure 8B). Moreover, the Bcl-2 motif identified in E-cadherin domain 4 is buried and thus not accessible to bulky residues of Bcl-2 structure (PDB ID: 2XA0). On the contrary, the motif found in domain 3 lies on the surface of the protein. Thus, while E-cadherin and Bcl-2 interaction with galectin-7 are similar, motif identified on domain 3 is a better candidate. Furthermore, a series of surface residues of E-cadherin, important for interaction between galectin-7 and E-cadherin, can be successfully structurally superposed on Bcl-2, confirming a similar interaction mode (figure 8B). These findings suggest that E-cadherin domain 3 is the most probable domain able to interact with galectin-7 which we confirmed by docking simulations. Furthermore, model of ectodomain 3 has good energy values all along the sequence and the motif of interest forms a β hairpin offering a great surface for a possible interaction.

Molecular docking experiments have led to an interesting pose that supports galectin 7/E-cadherin interaction. The best pose obtained is supported both in terms of energy (total docking score = 3812) and in terms of the statistical distribution of score values for the best first thousand poses obtained (Z-score = 5.20). This interaction model also corresponds to a region where the density of interaction in terms of pose is the highest (figure S5B) suggesting that the physico-chemical characteristics of the regions involved in the interaction are favorable and converge towards the optimal pose obtained. Moreover, the most interacting residues of E-cadherin in the top 1,000 docking poses include the ones composing the sequence motif identified previously. In addition, these residues were also at the interface in the best pose (Supp. Fig. 3). All these results greatly support the complex model.

We observed that E-cadherin extracellular domain 3 interacts with galectin-7 at the level of its dimerization interface (figure 8C, figure 8D and figure 8E). This region is found twice in galectin-7 and adopts a symmetrical arrangement which allows the binding of two E-cadherin molecules on both sides of galectin-7 (figure 8C, figure 8D and figure 8E). The binding of the two E-cadherin molecules is made in a symmetrical manner when docking first pose is considered. This pose is the most relevant, with a docking score highly superior to others (figure S5D). Thus both CRD domain of each galectin-7 are free to interact to sugar. The computation of solvent accessibility of galectin-7 CRD with and without the two E-cadherin molecules docked confirms that there are no significant differences (Table 1). It confirms the glycosylation-independent interaction mode of galectin-7 to E-cadherin. Finally, it also supports the capacity of galectin-7 to bind E-cadherin and EGFR through sugars at the same time. Indeed, EGFR adopts a particularly complex and flexible conformation, and can be observed as a monomer and a dimer which have a particular impact for EGFR function (Arkhipov et al., 2013). Thus, such interaction between E-cadherin and EGFR through galectin-7 would modify degree of freedom of EGFR and therefore impact its dynamic and function as it has been previously demonstrated in molecular dynamic simulation inter alia (Arkhipov et al., 2013). Also, even if it was not resolved in experimental structures, EGFR conformation is modulated by the binding of more than 10 long N-glycans. This type of sugars have very diverse lengths and molecular weights. Here, without experimental information about these sugars, a precise atomistic model cannot be proposed. However, galectin-7/E-cadherin interaction region is near 100 Å away from the membrane, a distance compatible with the apex part of EGFR. Considering that N-glycans glycosylating human EGFR can have very long chains, one can imagine a long-range interaction (< 40 nm as indicated by PLA) mediated by sugar chain. To strengthen this hypothesis, we further tested *in vitro* binding assays with purified proteins. Recombinant human galectin-7 (rGal7) was incubated with a recombinant chimeric E-cadherin containing the E-cadherin ectodomain (Asp155-Ile707) fused to a C-terminal 6-Histidine tag (E-cad-His) and with the EGFR ectodomain (Met1-Ser645) fused to a Fc fragment (Human IgG1-Fc (Pro100-Lys330) (EGFR-Fc). Experiment was conducted with protein G sepharose in such a way that only EGFR-Fc can bind to the beads. Strikingly, galectin-7 and E-cadherin were precipitated by EGFR extracellular domain (figure 8F). Interestingly, mutated galectin-7 with the R74S substitution in its CRD domain faintly precipitated *in vitro* with recombinant EGFR-Fc and E-cadherin. These results demonstrate a direct interaction between these three proteins reinforcing the concept of a tripartite complex.

**Table 1:** Relative solvent accessibility in of CRD domains of galectin-7 (defined by residues S1-P10, S30-P76 and L120-V127) with and without two E-cadherin molecules according to Naccess.

|  | Galectin-7 CRD Chain A | Galectin-7 CRD Chain B |
| --- | --- | --- |
| Without e-cadherin (Å<sup>2</sup>) | 2070.4 | 2121.0 |
| With 2 EC3 (Å<sup>2</sup>) | 1982.3 | 1986.4 |
| Difference (%) | 4.4 | 6.8 |

## DISCUSSION

From this study, we first pointed out that galectin-7 governs EGFR functions. We also highlighted a new link between EGFR and E-cadherin, galectin-7 not only bridges these two molecules but also regulates E-cadherin dynamics in response to EGF.

EGFR is a major actor in essential cellular processes such as differentiation, proliferation and cell migration. Its aberrant activity in numerous pathologies and notably in cancers underlies our need to understand the complex regulation of EGFR activity and downstream signaling events. Strikingly galectin-7 is known to play a critical role functioning as a regulator of keratinocyte proliferation and migration, as well as to maintain and restore epidermal homeostasis thus we address a new role of galectin-7 in EGFR regulation and function. Interestingly in this study we have highlighted galectin-7 as a new molecular partner for EGFR as we have pointed out that galectin-7 CRD domain can directly bind to the extracellular domain of EGFR *via* its glycosylation motifs. Moreover, as a consequence, we unravel that impairment of this interaction strongly increases EGFR phosphorylation and its main downstream pathways Akt, Erk and STAT3 inducing their over-activation. Interestingly our results reveal similarities with those obtained by Amaddi et al on flotillin. However membrane microdomain-associated flotillin proteins knockdown (flotillin-1/reggie-2) results in reduced EGF-induced phosphorylation of EGFR and in reduced activation of the downstream MAPK and Akt signaling (Amaddii et al., 2012). Hence galectin-7 and flotillin could have antagonistic role in the regulation of EGFR signaling pathways.

Signal transducing molecules have been shown to affect membrane trafficking but there are evidences that this regulation can be both ways, meaning that membrane trafficking can also regulate signal transduction events (Vieira et al., 1996). Activated EGFR is internalized and sorted at the early endosome. The fate of the receptor has important consequences for biological cell outputs, with the recycling pathway favoring cell proliferation, although the degradative pathway correlates with normal cellular homeostasis. Exposure to stress leads to the removal of the receptor from the cell surface, and this has been proposed to potentiate cell death. Conversely, stress-activated receptor might also be temporary or reversibly removed from the membrane, thereby promoting cell survival and/or proliferation (Tomas et al., 2014). This cell response to stress must be strongly considered in the case of the regulation by galectin-7 as the expression of galectins has been repeatedly related to stress situation (Advedissian et al., 2017a; Rabinovich and Toscano, 2009). Here we observe that galectin-7 depletion favors EGFR phosphorylation and ubiquitination. These early activation modification motifs on the membrane EGFR have been implicated in both endocytosis and degradation during EGF treatment. Nevertheless our results reveal that EGFR doesn’t seem to be more degraded in cells deprived of galectin-7 but on the contrary that a large part of the endocytosed EGFR is recycled to the plasma membrane, probably sustaining the higher and persistent level of activation of the EGFR pathways. Thus galectin-7 appears to be implicated in EGFR retention at the plasma membrane and in guiding EGFR to recycling pathway. In our experiments, loss of galectin-7 resulted in enhanced EGFR activation and recycling but also as a consequence in increased cell proliferation and migration. Such derailed endocytosis and recycling of cell-surface proteins, including EGFR/RTK, along with disturbed downstream signaling has been implicated in multiple cancers (Mosesson et al., 2008) and reinforced by our results on cell proliferation and migration.

Interestingly, the fate of endocytosed EGFR has been shown to be dependent of the nature of its ligand association. Indeed in the case of a high concentration of ligand, EGF mainly induced the major part of endocytosed EGFR to be degraded. On the contrary association of TGFα with EGFR led to a high proportion of recycling. These alternative trafficking of receptor is not due to differences in affinity but has been attributed to a quicker detachment of TGFα than EGF and at a higher pH (Ebner and Derynck, 1991). Yet the endosomal compartments are known to become more and more acidic when evolving to the lysosomal pathway thus TGFα detached probably earlier. Hence our results would suggest that galectin-7 by binding to EGFR stabilize the EGF bound receptor. In this way we showed in wild-type HaCaT cells the co-endocytosis of galectin-7 with EGFR suggesting that this interaction could stabilize the EGF/EGFR complex through the early step of endocytosis.

Besides, it has been published that N-linked glycosylation can define the individual properties of extracellular and membrane-associated proteins and that modifications of these glycosylations can alter constitutive cell proliferation and trigger epithelial-to-mesenchymal transition (Manwar Hussain et al., 2018). Interestingly it has been pointed out that inappropriate N-glycosylation often results in malfunction of EGFR such as in lung and breast cancers. The authors suggest that the allosteric organization of EGFR Tyrosine Kinase is dependent on extracellular N-glycosylation events and that EGFR functions would be linked to the N-glycosylation status of EGFR. Hence aberrant glycosylation would favor the constitutive activation of EGFR, conducting to proliferation and invasiveness. Hence the key role of glycosylation motifs in the control of endocytic pathways of decorated membrane protein has been recently reviewed with a special emphasis on the control of this phenomenon by galectins (Johannes and Billet, 2020). This role can be strengthened at the light of our results. Indeed, our study with the proposed model lead to the hypothesis that pathologic glycosylation of EGFR extracellular domain impairing galectin 7 binding would disrupt its control on proliferation and migration. It has been shown that several factors are able to regulate EGFR, even directly at the plasma membrane such as caveolin-1 (Tomas et al., 2014). We thus studied caveolin-1 which is a scaffolding molecule for several signaling proteins including EGFR. We observed in figure S5 that the recruitment of caveolin-1 is diminished in absence of galectin-7 and that caveolin-1 is less recruited in absence or in presence of exogenous EGF. These results are in accordance with the fact that phosphorylated EGFR is known to be excluded from caveolae (Abulrob et al., 2004). These results are also in concordance with the fact that chronic EGF treatment resulted in transcriptional downregulation of caveolin-1 with downregulation of E-cadherin expression (Lu et al., 2003). Indeed, many studies suggest that EGFR is involved in complexes with E-cadherin and catenins (Hoschuetzky et al., 1994; Pece and Gutkind, 2000). These studies demonstrate that while EGFR activation can disrupt E-cadherin function, E-cadherin can reversely antagonize EGFR activity (Hazan and Norton, 1998; Fedor-Chaiken et al., 2003). Taken together, these results indicate a dynamic relationship between EGFR and E-cadherin that regulates the function of both molecules.

We and others have previously published that a direct binding of galectin-7 can modulate the function or localization of its bound partners. For example, a fraction of galectin-7 is constitutively localized at mitochondria in a Bcl-2-dependent manner and sensitizes the mitochondria to the apoptotic signal (Villeneuve et al., 2011). We have also documented that galectin-7 is a direct partner of E-cadherin. Most noticeably this interaction has been linked to a sugar independent binding between galectin-7 and E-cadherin extracellular domain. In the present study, we show that galectin-7 also binds to EGFR through its extracellular domain in a carbohydrate dependent manner. Both these interactions regulate their plasma membrane dynamics and fate. In this study, we propose an *in silico* model of the bond between EGFR and E-cadherin through galectin-7. This robust model leads us to hypothesize that galectin-7 would limit the degree of freedom of EGFR which is otherwise a very flexible molecule rendering its endocytosis less efficient and possibly modifying its interaction with its ligand. The increase of stability at the plasma membrane that we observe in presence of galectin-7 for both EGFR and E-cadherin could be due to the formation of the whole three molecular complex that we described and fits well with all of our results. Thus, we reveal here for the first time that in keratinocytes galectin-7, can bind to both membrane proteins, EGFR and E-cadherin, sustaining at the molecular level their inter-regulation. The self-renewing epidermis is controlled by a balance between proliferation in the basal layer and a tightly controlled differentiation program with cells undergoing transcriptional and cell shape changes to form the distinct suprabasal layers. *In vitro* studies show that activation of EGFR plays an important role in reepithelialization by increasing keratinocyte proliferation and cell migration in acute wounds (Barrientos et al., 2008). Furthermore, it has been recently showed that E-cadherin is a master regulator of junctional and cytoskeletal tissue polarity in stratifying epithelia regulating the suprabasal localization and activation status of the EGFR essential to facilitate the development of a functional epidermal barrier (Rübsam et al., 2017). Galectin-7 downregulation in stratified epithelia has also been reported to be associated to development of several cutaneous manifestations and disorders and even to esophageal dysfunction in systemic sclerosis patient (Saigusa et al., 2019). To fulfill our study, *in vivo* observations in galectin7−/− mice compared to wild-type mice pointed out defects in epidermis differentiation in absence of galectin-7, hence revealing a thickening of the epidermis accompanied with an accumulation of K14 positive cells. Hence galectin-7 would control this differentiating step by regulating EGFR functions thus preventing excessive proliferation, migration and tumorigenic development.

In summary, we were able to demonstrate that negative control of EGFR phosphorylation by galectin-7 regulates EGFR activation and retention at the plasma membrane by changing its endocytosis. We were also able to demonstrate that galectin-7 bridges EGFR and E-cadherin in a tripartite molecular interaction and pointed out that galectin-7 controls the regulation of E-cadherin in response to EGF. As EGFR and E-cadherin are closely related and regulated we predicted a mechanism in which galectin-7 would be a major actor of epidermis differentiation through the regulation of the balance between EGFR and E-cadherin. These results provide unique mechanistic insight into how EGFR and E-cadherin would be interacting which is essential for the understanding of epithelial homeostasis and communication.

---

Figure S1: A/Representative immunoblots showing galectin-7 extinction in ShGal7 clones. B/ Absence of galectin-7 induces a downregulation of K10. Level of K14 and K10 transcripts have been compared in HaCaT cell and in ShGal7 clones after cell culture reached confluence. The transcript levels have been quantified from three independent quantitative PCRs. The p-value has been calculated by an ANOVA statistical test. C/ Level of K10 transcript has been compared in HaCaT cell and in ShGal7 clones in presence or in absence of EGF treatment at 100ng/mL for 16 hrs in HaCaT cells 0.5mM rGal7 has been added for 16hrs for one condition. Transcript levels have been quantified from three independent quantitative PCRs. The p-value has been calculated by an ANOVA statistical test.

Figure S2: Alteration of EGFR downstream pathways by galectin-7 deficient cells. A/ Immunoblots were probed for Phospho Src (Y416), Total Src. Gefitinib has been added at 15μM. Quantification of the blots are representative of at least 3 independant expériments. * p<0,1 **p<0,05 ***p<0,001 B/ Immunoblots were probed Phospho STAT3 and total STAT3. Gefitinib has been added at 15μM. Quantification of the blots are representative of at least 3 independant expériments. * p<0,1 **p<0,05 ***p<0,001

Figure S3: E-cadherin and EGFR interact in HaCaT cells. Co-immunoprecipitation experiments were realized with cell extract of a confluent cell monolayer and indicate that E-cadherin and EGFR interact in HaCaT cells. Image shown are representative of three independent experiments.

Figure S4: A/ Local Smith & Waterman alignments between Bcl-2 and E-cadherin a) ectodomain 3 (EC3) et b) ectodomain 4 (EC4). B/ Centers of gravity (as spheres) of E-cadherin top 1.000 docking poses around galectin-7 (in cartoon). Diameter and color of the spheres are proportional to docking energies. C/ Number of times an E-cadherin ectodomain 3 residue is found at the interface with galectin-7. D/ Scores computed by MEGADOCK default scoring function for the top 1.000 poses.

Figure S5: Caveolin-1 and EGFR are less interacting in absence of galectin-7. Cells have been treated or not with EGF at 100ng/mL for indicated time and then lysed. Immunoprecipitation of EGFR have been performed and western blot have been conducted. Blots have been probed with EGFR and caveolin-1 antibodies. Histograms represent caveolin-1 quantification normalized on HaCaT cells. Recombinant galectin-7 have been used at a concentration of 0.5mM for indicated time.

## Supplementary materials

| **Antibody** | **provider** | **reference** |
| --- | --- | --- |
| Galectin-7 | Abcam | ab10482 |
| EGFR | Abcam | ab52894 |
| P-EGFR | Abcam | ab40815 |
| P-Akt | Santa-cruz | sc-7985 |
| Akt | Cell signalling | 4691 |
| Erk | Santa-cruz | sc-154 |
| P-ERK | Santa-cruz | sc-7383 |
| P-Src | Cell signalling | 6943 |
| Src | Cell signalling | 2109 |
| STAT3 | Santa cruz | sc-293151 |
| P-STAT3 | Santa cruz | sc-8059 |
| Caveolin-1 | Cell signalling | 3267 |
| CD63 | Ancell | 215-020 |
| Lamp-1 | Santa-cruz | sc-20011 |
| E-cadherin | BD Biosciences | 610181 |
| GAPDH | Thermo Scientific | MA5-15738 |
| Anti-rabbit HRP | GE Healthcare | NA934V |
| Anti-mouse HRP | Sigma | A9044 |
| Anti-rabbit alexa 488 | Invitrogen | A11034 |
| Anti-mouse alexa 488 | Invitrogen | A11001 |
| Anti-rabbit alexa 568 | Invitrogen | A11011 |
| Anti-mouse alexa 568 | Invitrogen | A11004 |
| Keratin-10 | Covance | MMS-159S |
| Keratin-14 | Covance | PRB-155P |
| Transferrin fluorescein | Rockland | 00090234 |

## ACKNOWLEDGMENTS

This work was supported by Gefluc. We are grateful to Yves St Pierre for providing recombinant galectin-7 protein and the R74S mutant and Christian Wunder for providing fluorescently labelled galectin-7. We thank Marine Alves, Elise Grelet, Alix de Maupéou d’Ableiges de Montbail, Camille Gonzales and Coraline Hautem for valuable technical assistance Finally, we acknowledge the Buffon Animal Facility and the ImagoSeine core facility of the Institut Jacques Monod, member of IBiSA and the France-BioImaging (ANR-10-INBS-04) infrastructure.

## AUTHORS CONTRIBUTIONS

Veronique Proux-Gillardeaux is involved in conceptualization, funding acquisition, project administration, investigation, validation, visualization and, writing original draft. Tamara Advedissian is involved in conceptualization, validation, visualization and writing – review & editing. Jean-Christophe Gelly is involved in conceptualization, formal analysis, visualization and writing – review & editing. Charlotte Perin is involved in formal analysis, visualization and writing – review & editing. Mireille Viguier is involved in conceptualization, funding acquisition, investigation, validation project administration and writing original draft. Frederique Deshayes is involved in conceptualization, funding acquisition, investigation, validation, visualization, project administration and writing original draft.

