## Supplemental data for "Identification of a new regulation pathway of EGFR and E-cadherin dynamics"

Figure S1A

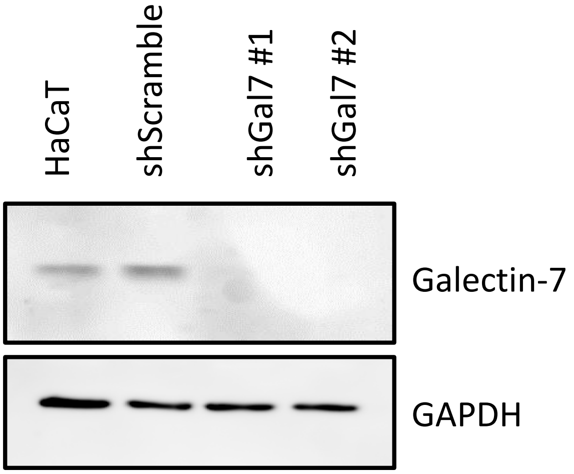


Figure S1B

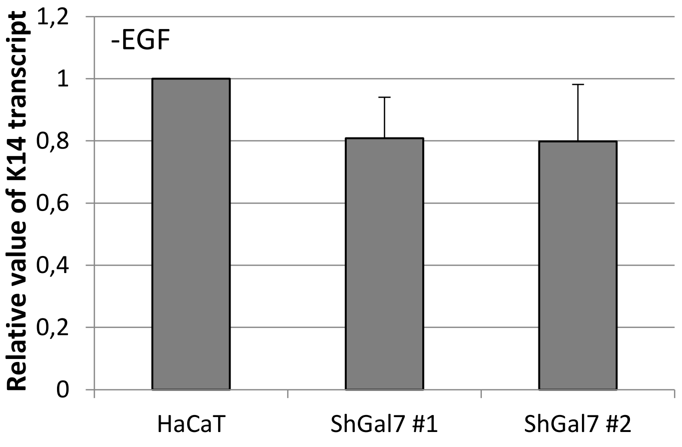


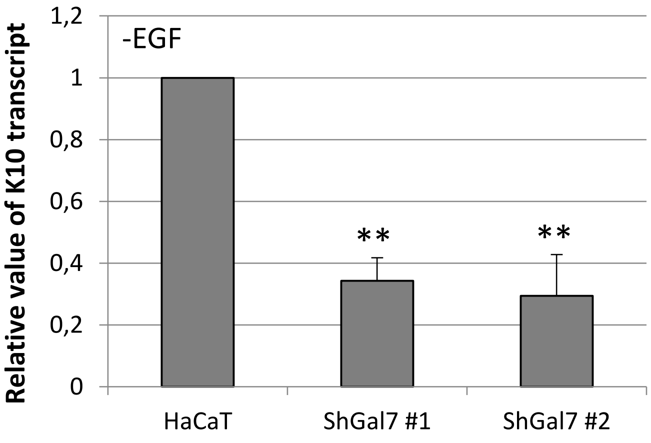

Figure S1C

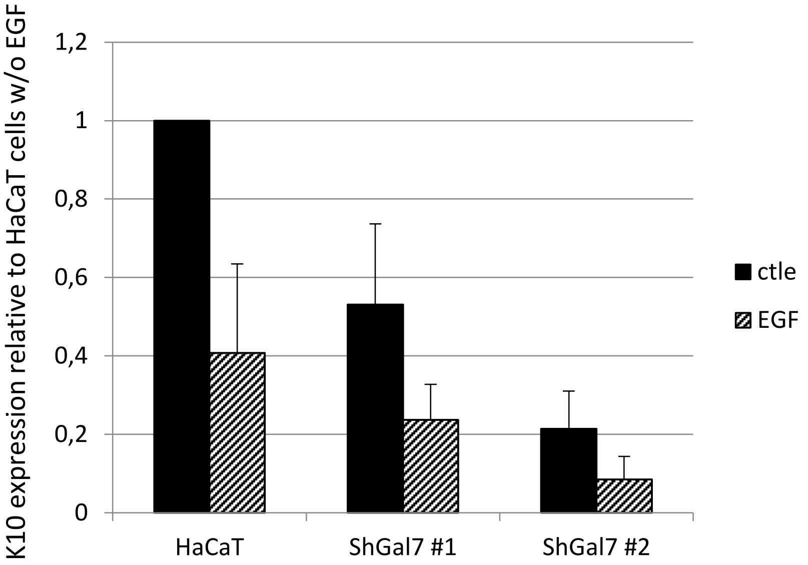
K10 expression relative to HaCaT cells w/o EGF
1,2
1
0,8
0,6
0,4
0,2
0
HaCaT
ShGal7 #1
ShGal7 #2
ctle
EGF

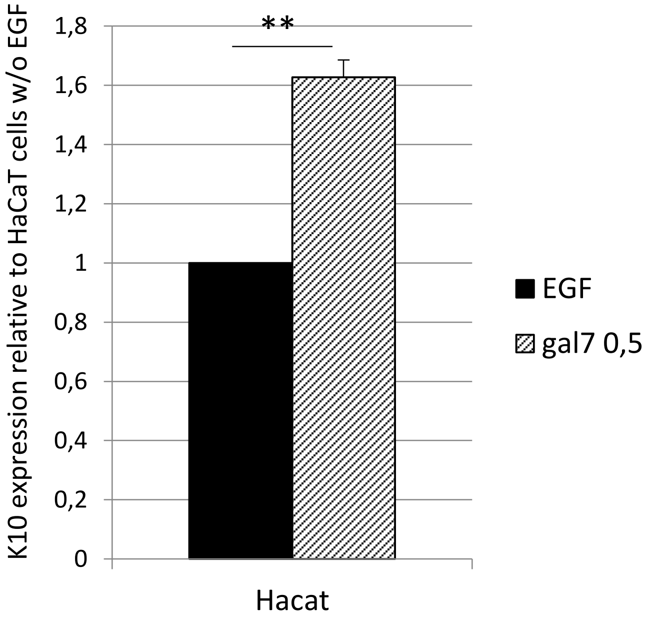
K10 expression relative to HaCaT cells w/o EGF
**
1,8
1,6
1,4
1,2
1
0,8
0,6
0,4
0,2
0
Hacat
EGF
gal7 0,5

figure S2A

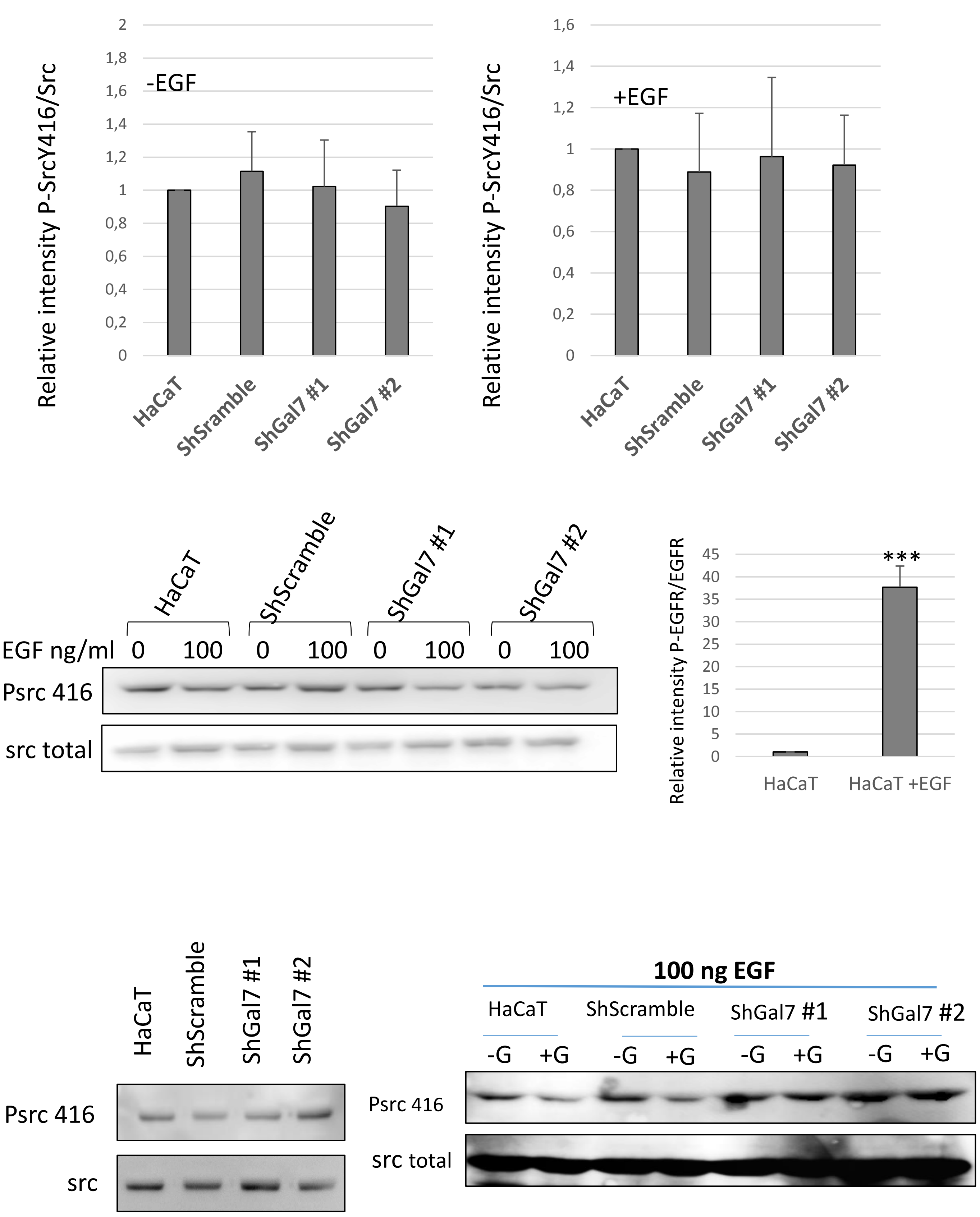

-EGF
Relative intensity P-SrcY416/Src
2
1,8
1,6
1,4
1,2
1
0,8
0,6
0,4
0,2
0
HaCaT
ShSramble
ShGal7 #1
ShGal7 #2
+EGF
Relative intensity P-SrcY416/Src
1,6
1,4
1,2
1
0,8
0,6
0,4
0,2
0
HaCaT
ShSramble
ShGal7 #1
ShGal7 #2
HaCaT
ShScramble
ShGal7 #1
ShGal7 #2
EGF ng/ml 0 100 0 100 0 100 0 100
Psrc 416
src total
Relative intensity P-EGFR/EGFR
45
40
35
30
25
20
15
10
5
0
***
HaCaT
HaCaT +EGF
HaCaT
ShScramble
ShGal7 #1
ShGal7 #2
Psrc 416
src
100 ng EGF
HaCaT
ShScramble
ShGal7 #1
ShGal7 #2
-G +G -G +G -G +G -G +G
Psrc 416
src total

figure S2B

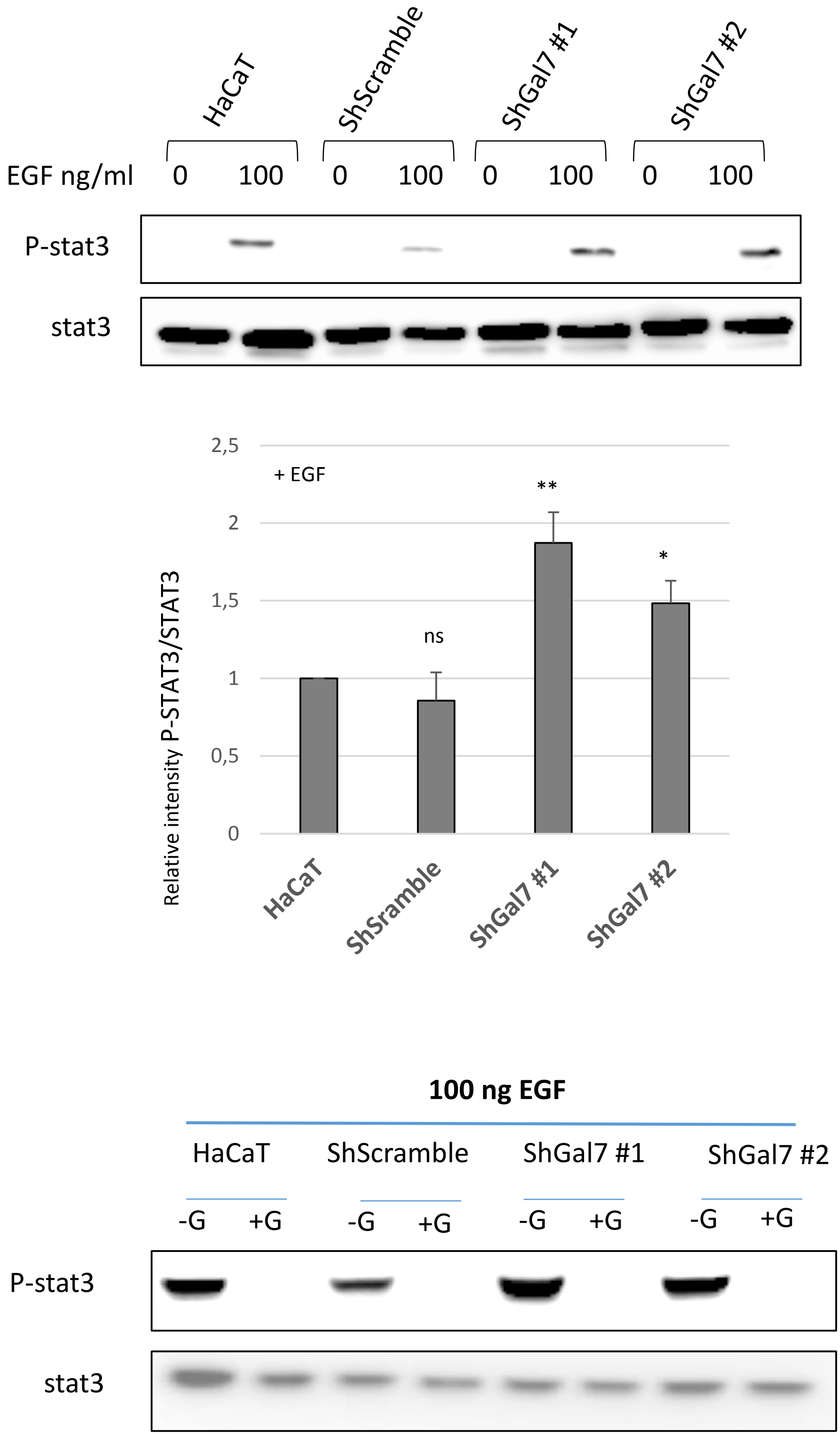
HaCaT
ShScramble
ShGal7 #1
ShGal7 #2
EGF ng/ml
0
100
0
100
0
100
0
100
P-stat3
stat3
2,5
2
1,5
1
0,5
0
+ EGF
**
*
ns
Relative intensity P-STAT3/STAT3
HaCaT
ShSramble
ShGal7 #1
ShGal7 #2
100 ng EGF
HaCaT
ShScramble
ShGal7 #1
ShGal7 #2
-G
+G
-G
+G
-G
+G
-G
+G
P-stat3
stat3

Figure S3

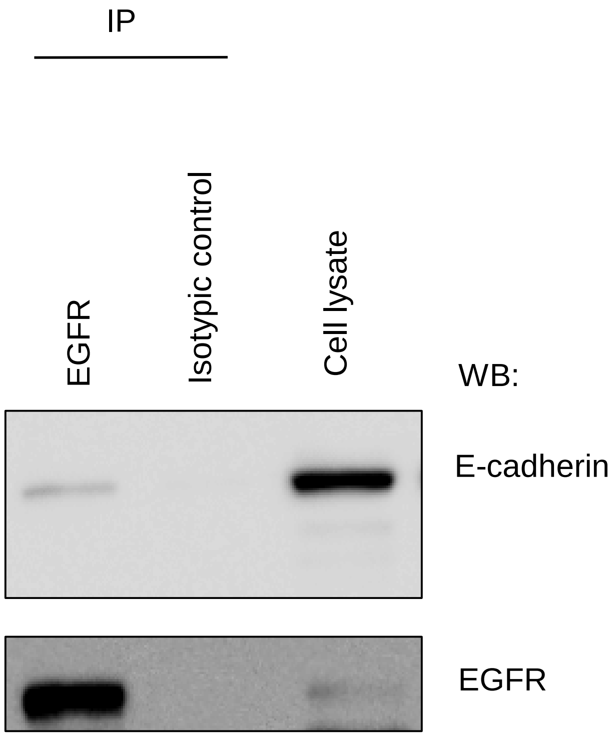

Figure S4A

**a**

```
EC3      NTPAWEAVY------TILNDDGG--QFVVTTNP
         |...|...|      |.:.|:||  .||....|
Bcl-2    NIALWMTEYLNRHLHTWIQDNGGWDAFVELYGP
```

**b**

```
EC4      ITSYTAQ-EPDTFMEQKITYRIWRDTANWLEINP--DTGAISTRAELDRE
         :|.:||: ...|.:|:     ::||..||..|..  :.|.:.....::||
Bcl-2    LTPFTARGRFATVVEE-----LFRDGVNWGRIVAFFEFGGVMCVESVNRE
```

Figure S4B

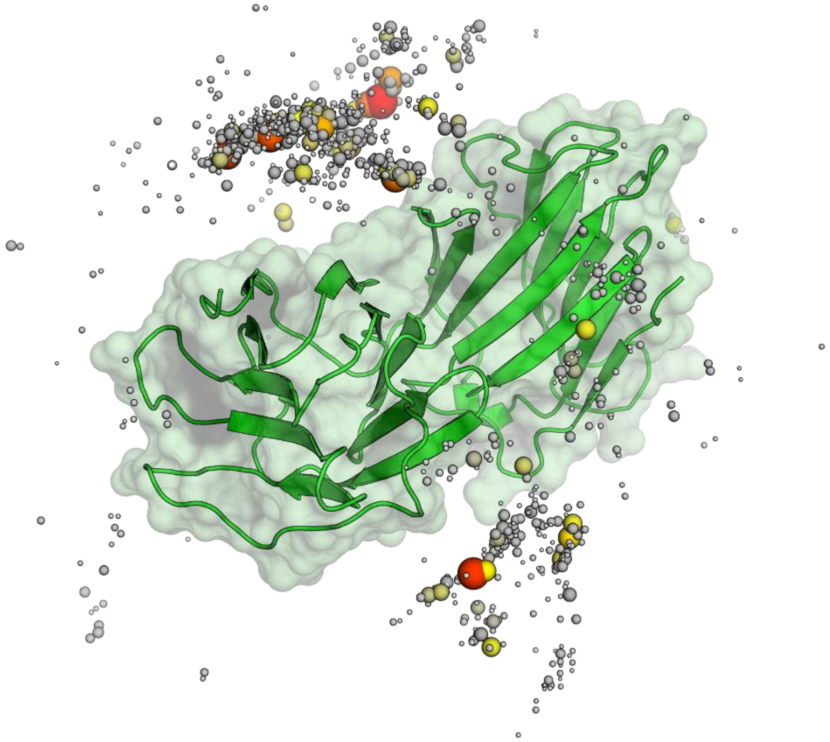

Figure S4C

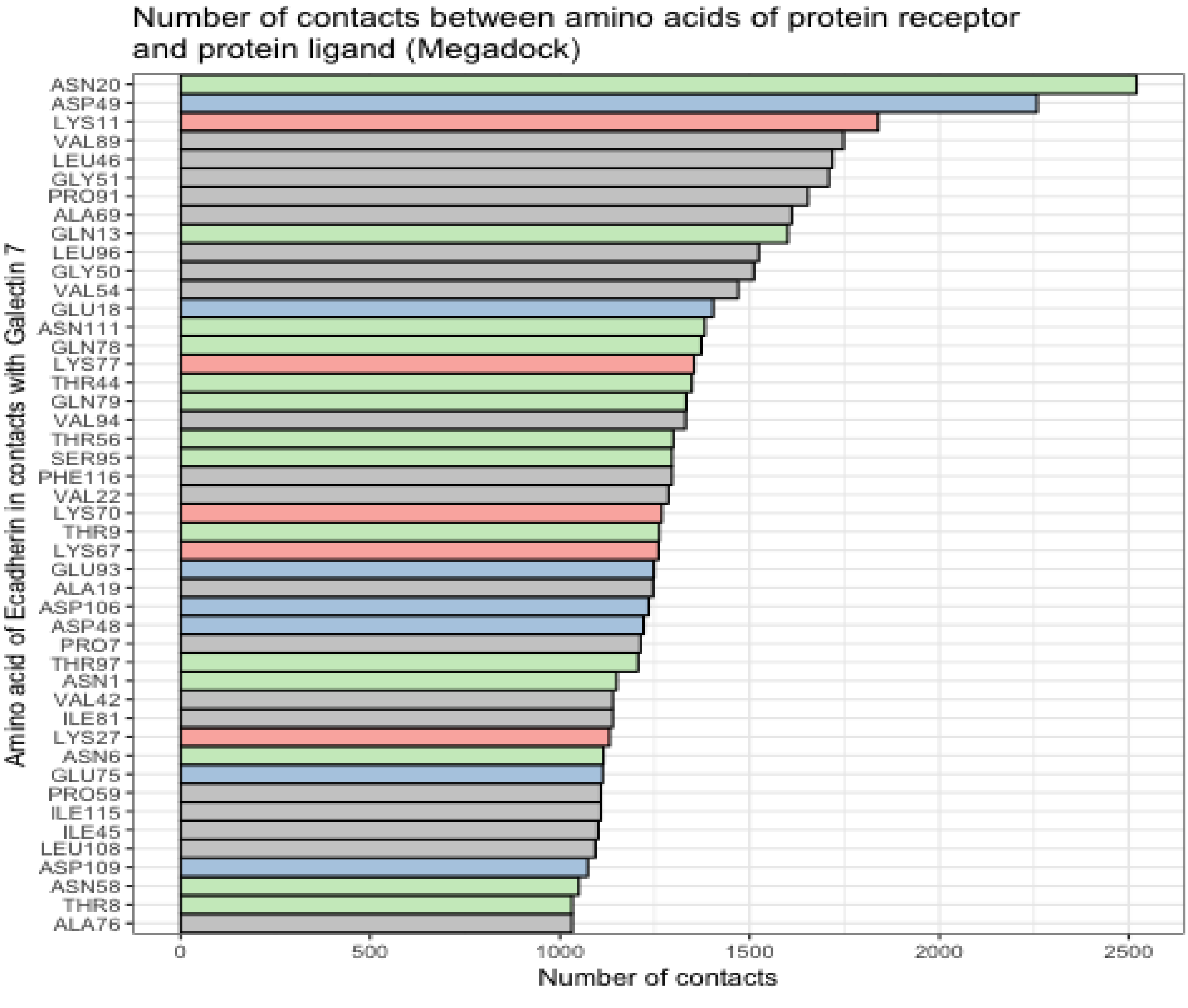

Number of contacts between amino acids of protein receptor and protein ligand (Megadock)
ASN20
ASP49
LYS11
VAL89
LEU46
GLY51
PRO91
ALA69
GLN13
LEU96
GLY50
VAL54
GLU18
ASN111
GLN78
LYS77
THR44
GLN79
VAL94
THR56
SER95
PHE116
VAL22
LYS70
THR9
LYS67
GLU93
ALA19
ASP106
ASP48
PRO7
THR97
ASN1
VAL42
ILE81
LYS27
ASN6
GLU75
PRO59
ILE115
ILE45
LEU108
ASP109
ASN58
THR8
ALA76
Amino acid of Ecadherin in contacts with Galectin 7
0
500
1000
1500
2000
2500
Number of contacts

Figure S4D

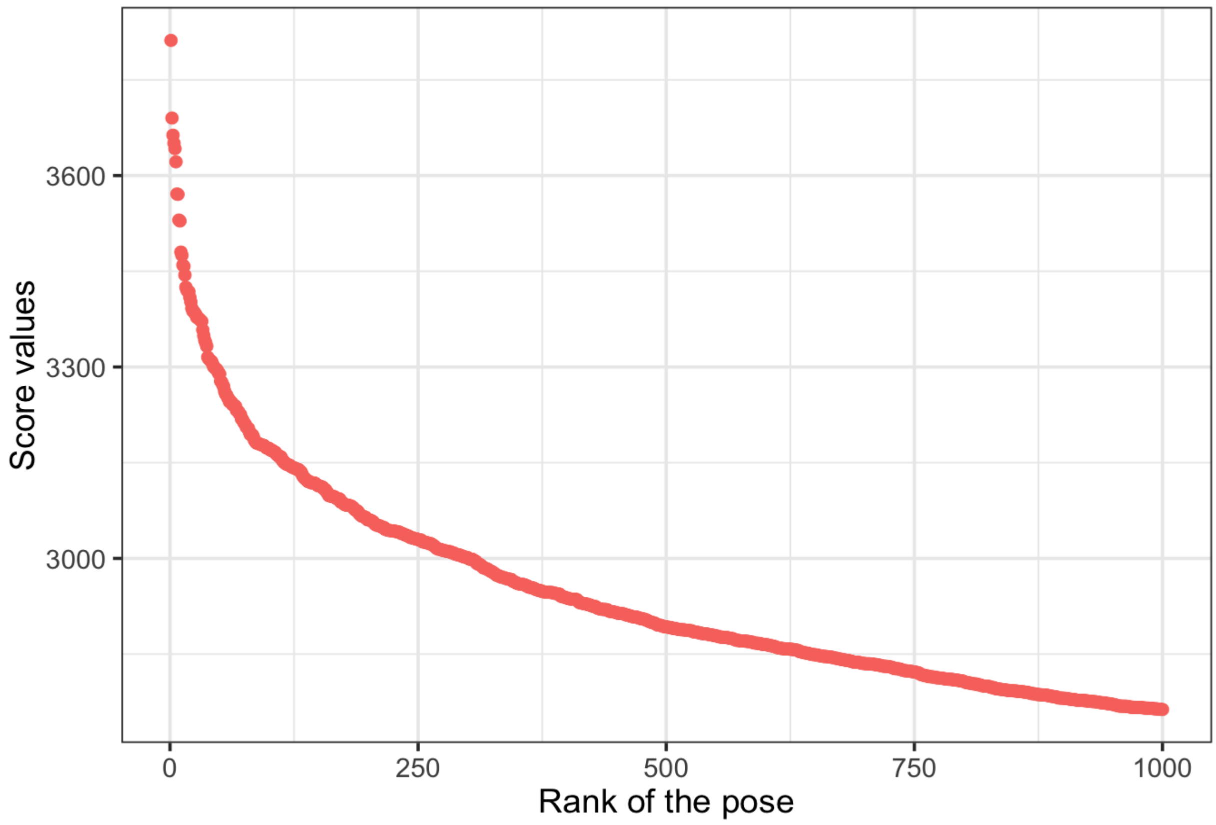
Score values
3600
3300
3000
0
250
500
750
1000
Rank of the pose

figure S5

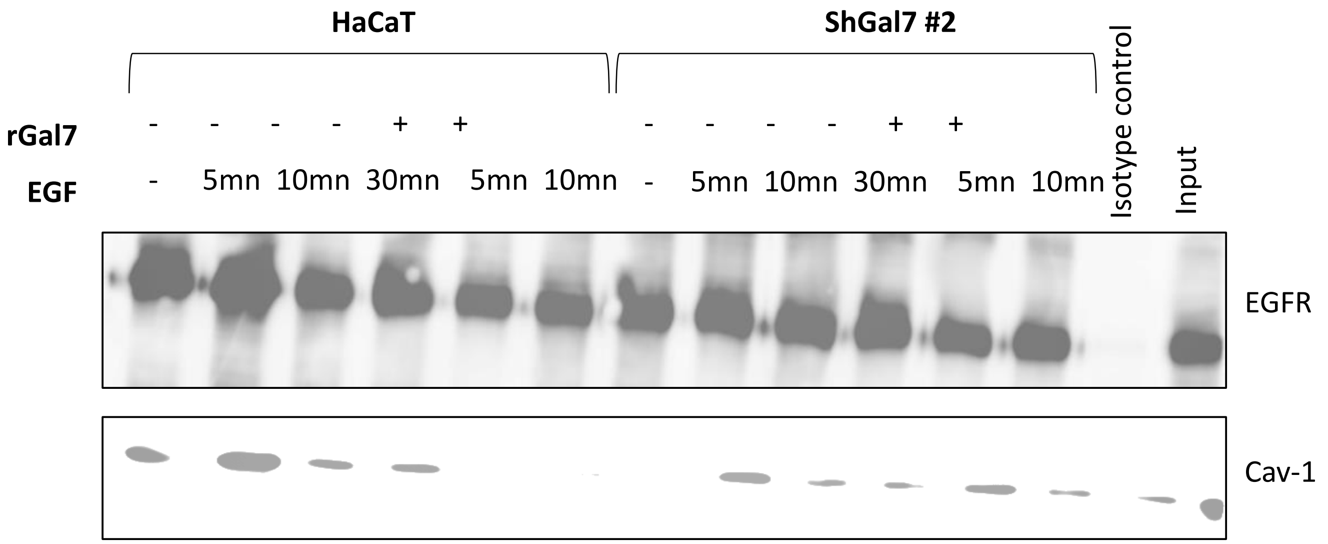
HaCaT
ShGal7 #2
rGal7
EGF
- - - - + +
- 5mn 10mn 30mn 5mn 10mn
- - - - + +
- 5mn 10mn 30mn 5mn 10mn
Isotype control
Input
EGFR
Cav-1

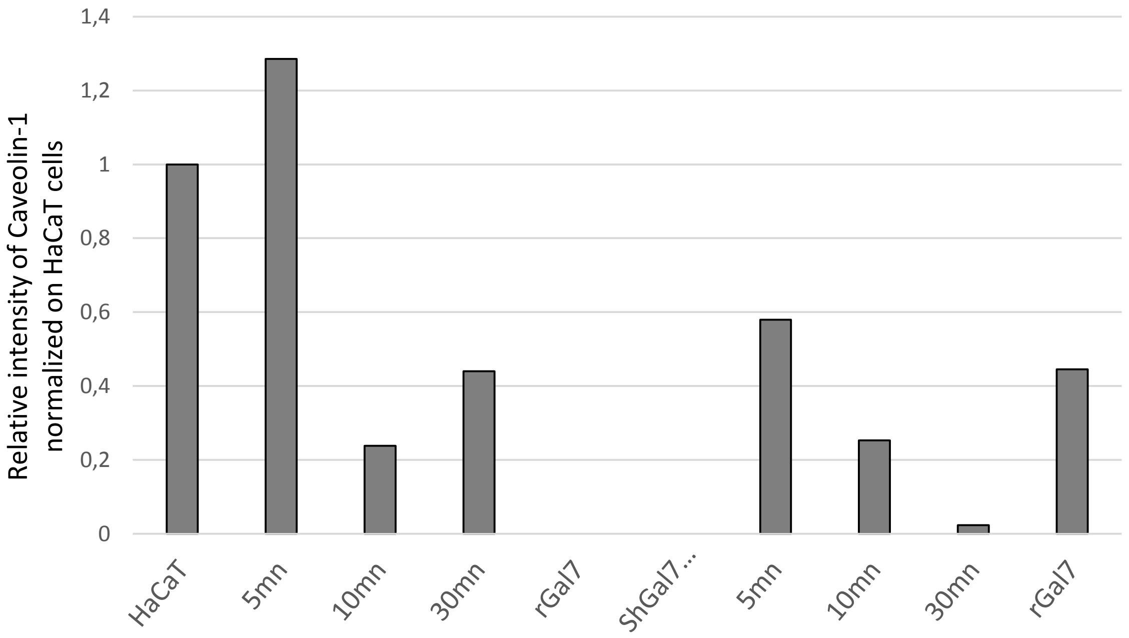
Relative intensity of Caveolin-1 normalized on HaCaT cells
1,4
1,2
1
0,8
0,6
0,4
0,2
0
HaCaT
5mn
10mn
30mn
rGal7
ShGal7 ...
5mn
10mn
30mn
rGal7
